# Submolecular video-imaging of the Smc5/6 complex topologically bound to DNA

**DOI:** 10.1101/2025.08.15.670630

**Authors:** Kenichi Umeda, Yumiko Kurokawa, Yasuto Murayama, Noriyuki Kodera

**Affiliations:** Nano Life Science Institute (WPI-NanoLSI), Kanazawa University, Kanazawa, Ishikawa, Japan; Department of Chromosome Science, National Institute of Genetics.; Department of Genetics, Graduate University for Advanced Studies (SOKENDAI), Mishima, Japan; PRESTO/JST, 4-1-8 Honcho, Kawaguchi, Saitama, Japan

## Abstract

Structural maintenance of chromosomes (SMC) complexes are ring-shaped ATPases that hold distinct DNA segments together to regulate various chromosome structures and functions. However, their dynamic mechanisms of DNA binding and processing remain poorly understood. Here, using high-speed atomic force microscopy, we directly visualized the dynamics of the Smc5/6 complex on DNA at submolecular resolution, resolving its individual domains. ATP-bound Smc5/6 stably binds DNA via the ATPase head domain, whereas following ATP hydrolysis, the DNA is entrapped within the SMC compartment and positioned around the opposite hinge domain. Furthermore, Smc5/6 topologically entraps two DNA segments together and stabilizes the DNA twist structures, promoting DNA compaction. Our findings provide a visual demonstration of how an SMC complex employs its ring architecture to facilitate distinct DNA binding modes.

## Introduction

Members of the structural maintenance of chromosomes (SMC) proteins, including eukaryotic cohesin (Smc1/3), condensin (Smc2/4) and the Smc5/6 complex, play fundamental roles in chromosome organization and segregation (*1*). SMC complexes are thought to fold genomic DNA by mediating physical interactions between two distinct chromosomal segments. SMC complexes are characterized by their ring-like architecture, which is composed of a pair of SMC ATPase subunits and a kleisin subunit that further interacts with various regulatory subunits. Studies on purified SMC complexes have revealed diverse DNA transactions driven by ATP binding and hydrolysis. Cohesin, in particular, is well known for its ability to topologically entrap DNA within its ring, which underpins the establishment of sister chromatid cohesion (*2*, *3*). In addition, all eukaryotic SMC complexes, including Smc5/6, have been shown to extrude DNA loops in a continuous manner (*4–7*). This unique function may underlie the role of cohesin in shaping interphase genome architecture and of condensin in driving chromosome compaction during mitosis. DNA loop extrusion by prokaryotic SMC complexes has also recently been reported (*8*). Moreover, cohesin and condensin have been shown to tether two separate DNA strands, offering an alternative molecular mechanism for establishing DNA–DNA interactions (*9–11*). However, it remains unclear how SMC complexes bind to DNA and exploit their ring-like architecture to carry out their functions, particularly from a dynamic structural perspective.

Smc5/6 is essential for accurate chromosome segregation during both mitosis and meiosis (*12–14*). This is primarily due to its role in eliminating harmful DNA structures that can hinder DNA replication, including repair intermediates of DNA damage through homologous recombination, stalled replication forks, DNA–RNA hybrids (R-loops), DNA supercoils, and DNA intertwined between sister chromatids (*15–19*). Purified Smc5/6 binds directly to DNA and stabilizes DNA supercoils in an ATP-dependent manner (*20*, *21*). Smc5/6 has been shown to exhibit a higher affinity for single-stranded and double-stranded DNA junctions formed during DNA replication and recombinational repair, suggesting that it can recognize higher-order DNA structures (*22*, *23*). In addition, like cohesin and condensin, Smc5/6 is capable of topological DNA entrapment (*24*). These unique DNA interactions appear to be directly linked to the functions of Smc5/6 in a wide range of chromosomal processes. Therefore, it is crucial to elucidate the structural dynamics of Smc5/6 on DNA in order to gain further molecular insights into how these complexes enable various DNA interactions.

Recent studies using single-molecule fluorescence imaging have revealed the molecular dynamics of Smc5/6 but lacked the structural details (*7*, *19*, *22*, *23*). Cryo-electron microscopy studies have provided atomic-level structures of Smc5/6, albeit with limited dynamic information (*25–28*). High-speed atomic force microscopy (HS-AFM) has rapidly developed as a unique approach to visualize the nanodynamics of biomolecules in an active state in real space under physiological buffer conditions (*29*, *30*). This approach has revealed the molecular behavior of SMC proteins including cohesin, condensin and the SMC-related MRN DNA repair complex (*31–33*). By contrast, the dynamics of DNA-bound SMC proteins have not been adequately addressed, as the previous studies have primarily used a conventional dry AFM or samples strongly adsorbed electrostatically to charged AFM stages (*32*, *34*, *35*).

Here we visualized the dynamics of *Saccharomyces cerevisiae* Smc5/6 topologically bound to DNA using HS-AFM combined with a mica-supported lipid bilayer system, which has been successfully applied to investigating the intrinsic nature of proteins bound to scaffold molecules such as DNA and cytoskeletons (*36–38*). Our observations provide deeper mechanistic insights into the ATP-driven functions of Smc5/6.

## Results

### The Smc5/6 core complex adopts both I- and O-shaped conformations

The Smc5/6 complex consists of eight subunits, including a pair of SMC subunits (Smc5 and Smc6) and six non-SMC element (NSE) subunits that form a stable dimer via hinge domains. The other ends of the extended SMC coiled-coil arms are ATPase heads that engage upon ATP binding. The Nse4 kleisin subunit bridges the two SMC heads through asymmetric interactions, forming a closed ring topology. Nse1 and Nse3 stably interact with Nse4, while Nse2 associates with Smc5 at the center of its coiled-coil arm. The core of Smc5/6 consists of these six subunits (hereafter referred to as the Smc5/6 hexamer) (Fig. 1A). Further interaction with the Nse5–Nse6 subcomplex, which regulates topological DNA loading of the core complex, results in the formation of an octameric complex (hereafter referred to as the Smc5/6 octamer).

**Figure 1.**
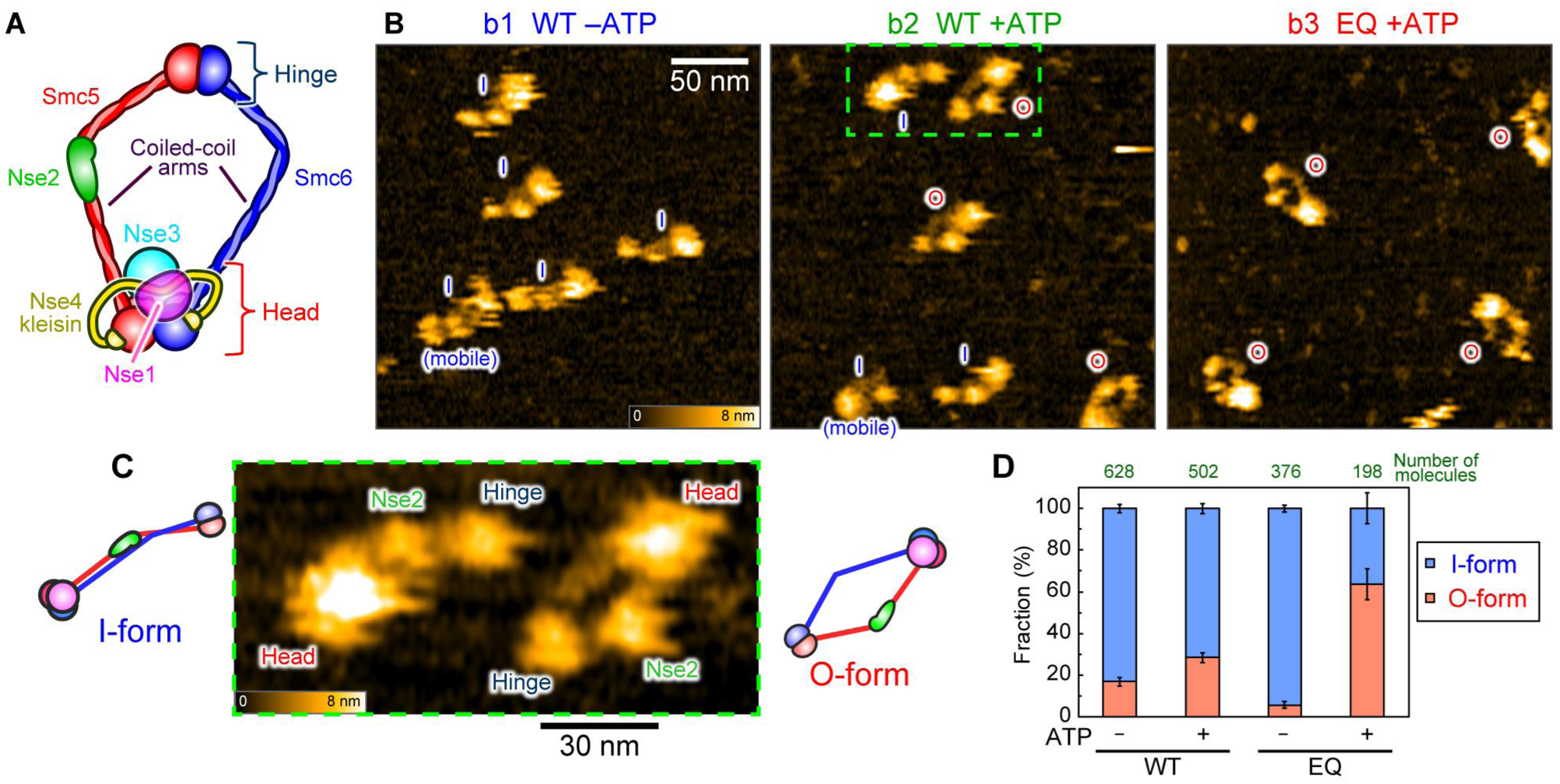
Structural evaluations of the budding yeast Smc5/6 hexamer complex on mica. (**A**). Schematic of the Smc5/6 hexamer. (**B**) Representative overview HS-AFM images of Smc5/6 hexamer molecules on a mica imaging stage. Wild-type (WT) hexamers without ATP (b1), with ATP (b2), and the EQ mutant molecules with ATP (b3). b1–b3: Scan area, 250 × 250 nm^2^ at 200 × 100 pixels^2^. Imaging rate, 1.5 s per frame. The molecules labeled “mobile” were identified as the I-form in adjacent frames but appear blurred here because all molecules are in motion. (**D**) Enlarged views of representative I-form and O-form wild-type hexamers (from b2). (**D**) Abundance ratios of different conformations calculated from individual molecules (obtained from three independent experiments), where error bars represent s.d..

We purified the budding yeast *Saccharomyces cerevisiae* Smc5/6 complex as both octameric and hexameric forms, which exhibited DNA-stimulated ATPase activity as previously reported (fig. S1) (*39–41*). These purified complexes were subjected to HS-AFM observation in solution on freshly cleaved bare mica, a common imaging stage for supporting biomolecules. A buffer solution containing a high concentration of salt (300 mM KOAc) was used to minimize the disruption of Smc5/6 during HS-AFM observation. We first observed several hexamer molecules simultaneously within the same viewing window (Fig. 1B). This allowed us to visualize two types of Smc5/6 hexamers; an arm-open, ring shape (O-form) and an arm-closed, extended shape (I-form) (Fig. 1C). We quantified the occurrence frequencies of these two structures based on the conformations observed within the initial ten frames after the molecules appeared in the AFM images. More than 80% of Smc5/6 hexamers were in the I-form in the absence of ATP, with a slight increase in the O-form molecules (∼28%) in the presence of ATP (Fig. 1D).

We also observed the EQ mutant hexamer, which is capable of ATP binding but defective in hydrolysis (Fig. S1B, D) (*40*, *41*). Approximately 60% of the EQ mutant complex displayed the O-form in the presence of ATP, whereas most of the molecules remained in the I-form without ATP (Fig. 1B, D). This suggests that the engagement of the ATPase heads, mediated by ATP binding, induces the conformational change to O-form of the Smc5/6 core complex. In addition to hinge and head regions, an additional globular mass was observed at the middle of one of the SMC arms, corresponding to Nse2, which is known to bind the Smc5 coiled-coil region (*42*). This characteristic domain thus allowed us to distinguish Smc5 from Smc6 (Fig. 1C). In contrast to the Smc5/6 hexamer, our attempt to visualize the octamer molecules failed because Nse5-Nse6 appeared to become dissociated from the Smc5/6 hexamer core complex upon adsorption onto mica (fig. S2).

### ATP-dependent conformational changes of the Smc5/6 core complex

The I-form observed in our HS-AFM analyses is reminiscent of the cryo-electron microscopy (cryo-EM) structure of the budding yeast Smc5/6 hexamer in the absence of ATP (Fig. 2A, B) (*25*). Accordingly, the pseudo-AFM image generated from the cryo-EM structure by AFM image simulation (Fig. 2C, wild-type, apo state, see Materials and Methods) was virtually identical to the experimentally observed Smc5/6 hexamer using HS-AFM (Fig. 2A), with similar domain-to-domain distances, including hinge to Nse2 and Nse2 to ATPase heads (15 nm and 23 nm for the experiment, 16 nm and 23 nm for the pseudo-AFM image respectively). In contrast to the apo state (I-form), the previously published cryo-EM structure of the ATP-bound state, in which the ATPase heads and Nse1/Nse3 clamp DNA, was confined to the ATPase head region (*26*). This prompted us to construct an atomic model of the O-form to simulate the structural transition from the apo state (I-form) to the ATP-bound state (O-form), guided by our HS-AFM observations of the O-form molecules (Fig. 2D) and previously published cryo-EM structures of both states (*25*, *26*).

**Figure 2.**
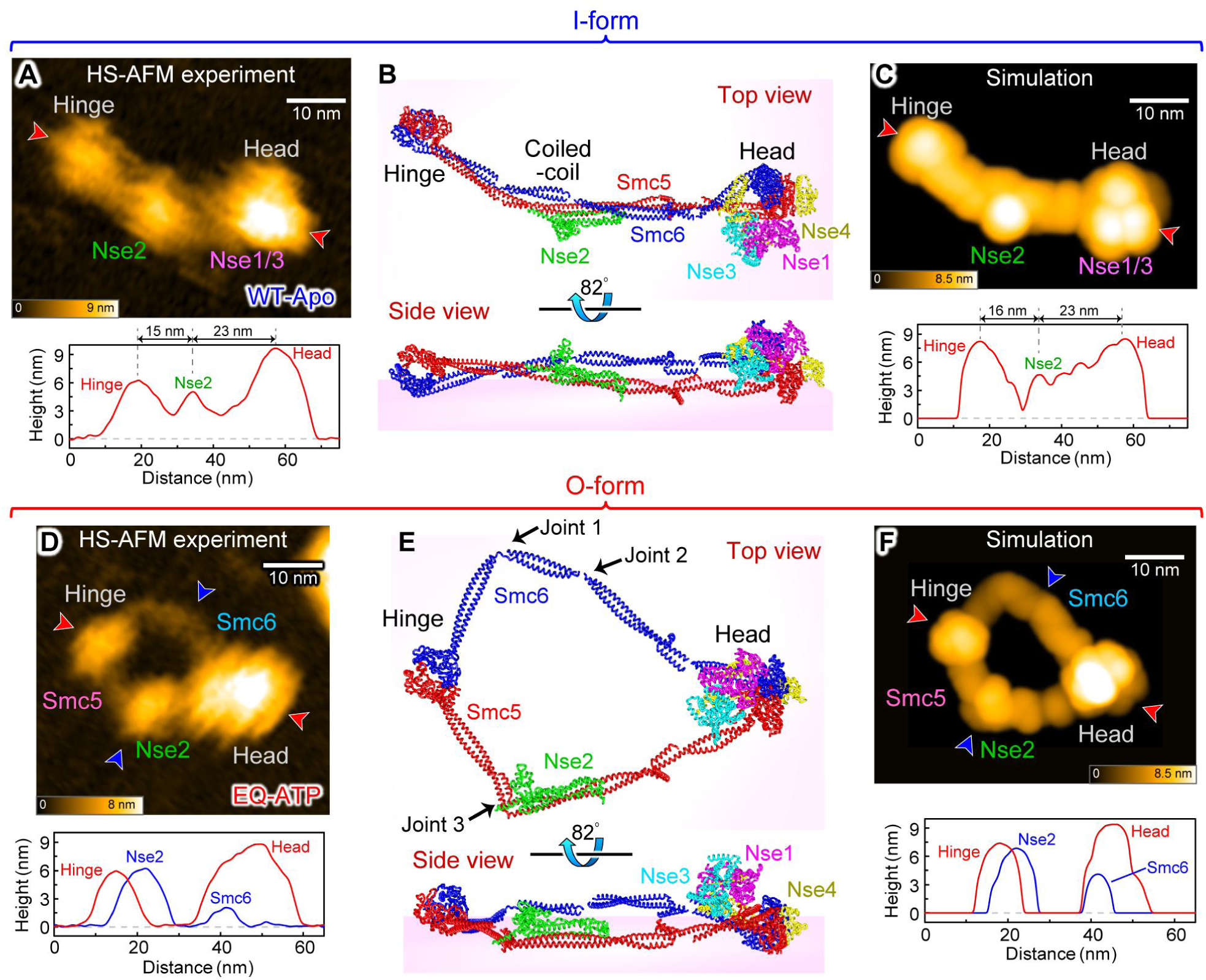
Molecular prediction of ATP-dependent conformational change of Smc5/6 hexamer. (**A**) A representative HS-AFM zoomed image of the I-form Smc5/6 hexamer (wild-type without ATP). Scan area, 100 × 100 nm^2^ at 160 × 80 pixels^2^, cropped to 73 × 56 nm². Imaging rate, 0.5 s per frame. (**B**) Atomic model of the budding yeast Smc5/6 hexamer in the apo state from its cryo-EM structure (EMD-13895, PDB-7QCD) (related to movie S1). (**C**) A pseudo-AFM image of the Smc5/6 hexamer in the apo state using the atomic model from **b**. In panels a and c, below the images, the corresponding line profiles of the molecular height across the two red arrowheads are presented. (**D**) A representative HS-AFM zoomed image of the O-form Smc5/6 hexamer (EQ mutant with ATP). Scan area, 100 × 100 nm^2^ at 160 × 80 pixels^2^, cropped to 55 × 55 nm². Imaging rate, 0.5 s per frame. (**E**) A predicted atomic model of Smc5/6 hexamer in the ATP-bound state (see fig. S3 and Materials and Methods) (related to movie S1). (**F**) A pseudo-AFM image of the Smc5/6 hexamer in the ATP-bound state using the atomic model from **e**. In panels d and f, below the images, the corresponding line profiles of the molecular height across the two red and blue arrowheads are presented, respectively.

Both the I- and O-form molecules observed by HS-AFM exhibited consistently extended SMC arms, leading us to speculate that the SMC arms can be considered as rigid chains. Therefore, we constructed a coarse-grained model based on a freely-jointed chain approach (*43*), using the published cryo-EM structure of the apo-state Smc5/6 (fig. S3, table S1, S2) (*25*). The SMC arms are composed of long antiparallel α-helices with several discontinuities (joints). Our HS-AFM observations of the O-form molecules showed that both SMC arms bent at obtuse angles near the middle. Consistent with this interpretation, Smc6 contains two joints where the two antiparallel peptide strands are disordered at the same positions in the middle of its arm. Smc5 also contains a similar joint, located near Nse2 on the hinge side. Therefore, we placed one and two flexible points on Smc5 (joint 3) and Smc6 (joint 1 and 2) arms, respectively (Fig. 2E). We also placed two flexible points at the bases of the arms, near the hinge domains. A harmonic oscillator model-type force was applied between the ATPase heads to recapitulate the ATP-dependent head engagement.

Next, we connected these flexible points with rigid chains to generate a simple framework. The I-form framework was converted into the O-form framework by applying virtual external pulling forces (Fig. 2E, fig. S3, movie S1), guided by the ATP-bound cryo-EM structure (*26*). The conversion to the O-form was only observed when the forces were applied to joint 1 and 3 in opposite directions (i.e., pulling apart). Finally, we generated the pseudo-AFM image based on the simulated O-form atomic model (Fig. 2F, see Materials and Methods). This modeling produced an image essentially identical to the experimentally observed Smc5/6 O-form. Together, our molecular simulation and HS-AFM analysis strongly suggest that Smc5/6 undergoes dynamic structural transitions between the I-form and O-form during ATP hydrolysis cycles.

To directly track the ATP-dependent conformational changes of the Smc5/6 hexamer, we further scanned these molecules over time within smaller scanning ranges. While the Smc5/6 hexamer primarily remained in the I-form without ATP, it exhibited transitions between I-form and O-form in the presence of ATP (Fig. 3A, movie S2). To quantify these conformational changes, we measured the arm–arm distances between Smc5 and Smc6 at the center of their arms in each time frame (Fig. 3B, C). The histograms derived from 10 molecules of Smc5/6 hexamers showed a large single peak in the absence of ATP, whereas two distinct peaks were observed in the presence of ATP. Gaussian fitting of the histograms was performed to calculate the mean arm–arm distances in each condition: 5 nm (I-form) in the absence of ATP, and 8.5 (I-form) and 19 nm (O-form) in the presence of ATP. Like the wild-type, the EQ mutant showed two distinct peaks in the histogram in the presence of ATP. However, in contrast to the wild-type, the majority (75%) of the EQ mutants were assigned to species with larger arm–arm distances with a mean distance of 18 nm (O-form) (Fig. 3B, C, movie S2). These results provide further support for the conclusion that ATP binding and subsequent hydrolysis regulate the conformational changes of the Smc5/6 core complex.

**Figure 3.**
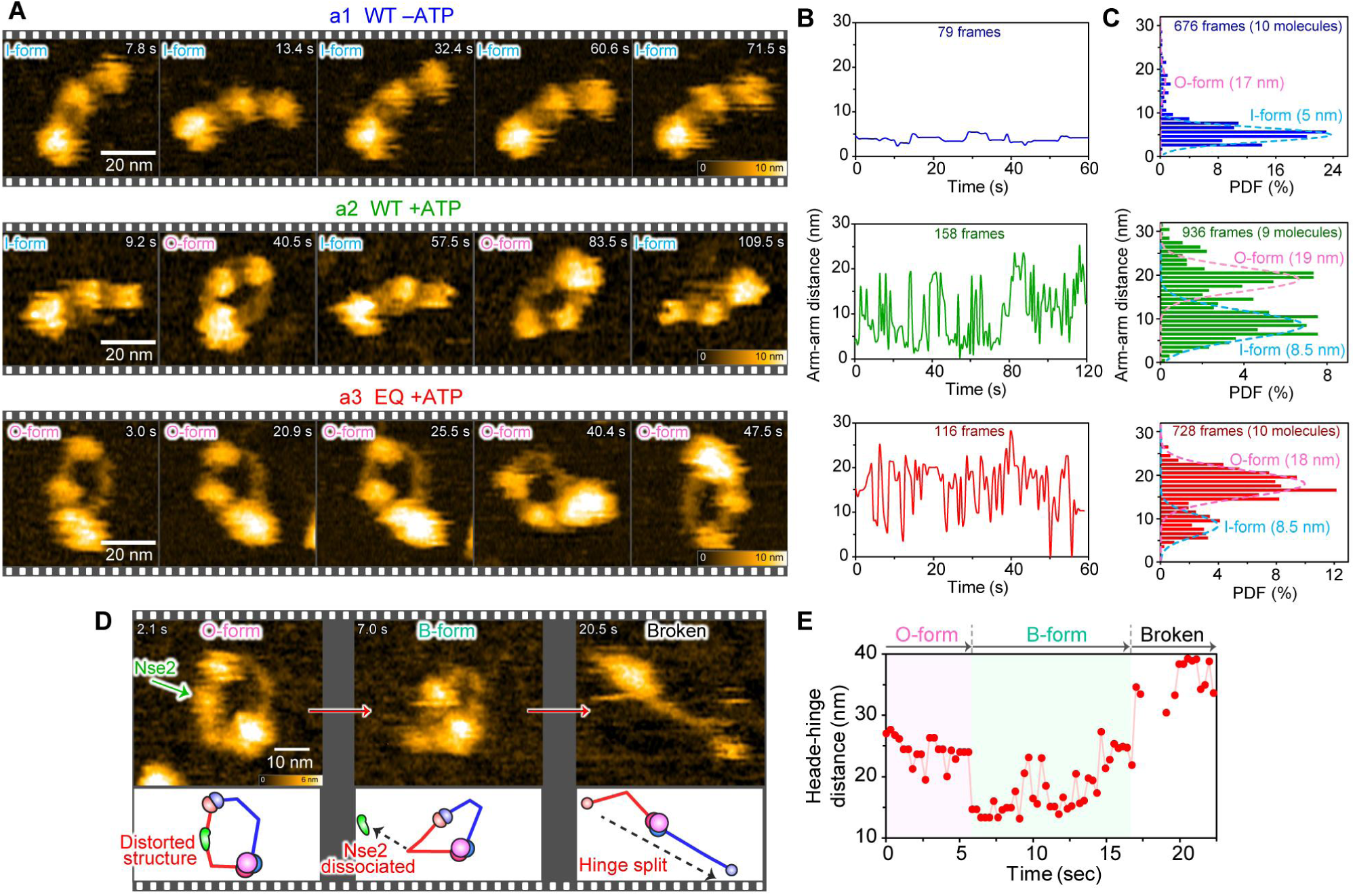
Real-time tracking of structural dynamics of Smc5/6 on mica. (**A**) Successive HS-AFM images of wild-type hexamers without ATP (a1), with ATP (a2), and the EQ mutant hexamers with ATP (a3) on a mica imaging stage (related to movie S2). a1 and a3: Scan area, 100 × 100 nm^2^ at 160 × 80 pixels^2^, cropped to 65 × 60 nm². Imaging rate, 0.5 s per frame. a2: Scan area, 150 × 150 nm^2^ at 160 × 80 pixels^2^, cropped to 75 × 63 nm². Imaging rate, 0.7 s per frame. (**B**) Real-time traces of arm−arm distance between Smc5 and Smc6, analyzed from the molecule shown in panel (**A**). (**C**) Distributions of arm−arm distances extracted from successive HS-AFM images under each condition. The total number of frames and analyzed molecules are indicated in each chart. The values in brackets next to each component label represent the fitted mean values of the Gaussian distributions. (**D**) Successive HS-AFM images of wild-type hexamer molecules undergoing Nse2 dissociation, observed in a buffer solution with a low KOAc concentration (75 mM) (related to movie S3). Scan area, 100 × 100 nm^2^ at 200 × 100 pixels^2^, cropped to 64 × 56 nm². Imaging rate, 0.3 s per frame. (**E**) Real-time trace of the distance between the hinge and head domains, shown in panel (**D**).

### Nse2 supports the extended state of the SMC arms

Although the Smc5/6 hexamer adopted two distinct shapes (I- and O-forms), its SMC arms remained extended and never folded back. This characteristic differs from cohesin and condensin, whose arms can sharply fold back at the “elbow”, displaying a ‘B’-form structure (*32*, *33*). Unlike cohesin and condensin, the Smc5/6 complex contains Nse2, which stably associates with the Smc5 arm (*42*), suggesting that Nse2 constrains the bending of the Smc5 arm. This notion is supported by the following two observations. First, in the O-forms of the EQ mutant, we observed that the Smc5 arm was constantly curved at the Nse2 binding region, whereas the Smc6 arm fluctuated due to repeated bending and stretching (fig. S4A). The histograms of the curvature angles at the Smc5 kink exhibited a single peak with a mean value of 50°, whereas Smc6 showed two distinct peaks at 6° and 40° (fig. S4B, C). These differences suggest that the Smc5 arm is stiffer and fixed at a specific angle, whereas the Smc6 arm is more flexible. Second, in a buffer solution with a low salt concentration (75 mM KOAc), we frequently observed the collapse of the Smc5/6 complex. During this process, the Smc5 arm exhibited sharp bending along with Smc6, which was associated with Nse2 dissociation, resulting in the formation of a ‘B’-like structure (Fig. 3D, E; movie S3). Together, these observations suggest that Nse2 acts like a cast, stabilizing the Smc5 arm angle.

### Observation of the Smc5/6 complex bound to DNA

To visualize DNA-bound Smc5/6 by HS-AFM, we first reconstituted the DNA loading of the Smc5/6 using previously established assays (Fig. 4A) (*3*). Since Nse5–Nse6 has been reported to be essential for the efficient topological DNA loading of Smc5/6(*24*), we used the Smc5/6 octamer for the reconstitution of DNA loading. Following incubation with circular DNA, Smc5/6 was retrieved by immunoprecipitation through a 3×Pk tag on Smc6, and the bound DNA was analysed by agarose gel electrophoresis after a high-salt wash. DNA recovery was observed when Smc5/6 was incubated in the presence of ATP, but not with ADP, cytidine triphosphate (CTP), or in the absence of nucleotides (Fig. 4B). Single-site restriction digestion of the captured DNA released it from the bead-bound Smc5/6, confirming the topological nature of Smc5/6 loading in the presence of ATP (fig. S5A). Similar results were observed with the EQ mutant Smc5/6 octamer, indicating that efficient ATP hydrolysis is not rate-limiting for topological Smc5/6 loading (fig. S5B). Consistent with this conclusion, the efficient DNA recovery with Smc5/6 was also observed with Smc5/6 in the presence of the non-hydrolysable ATP analogue ADP-AlF_x_ (Fig. 4B), which has been reported to promote topological DNA loading of cohesin (*44*, *45*). Although another analogue, ATP-γ-S, did not support DNA loading of Smc5/6, ATP-γ-S appears to be an imperfect ATP mimic for Smc5/6, similar to cohesin (*3*).

**Figure 4.**
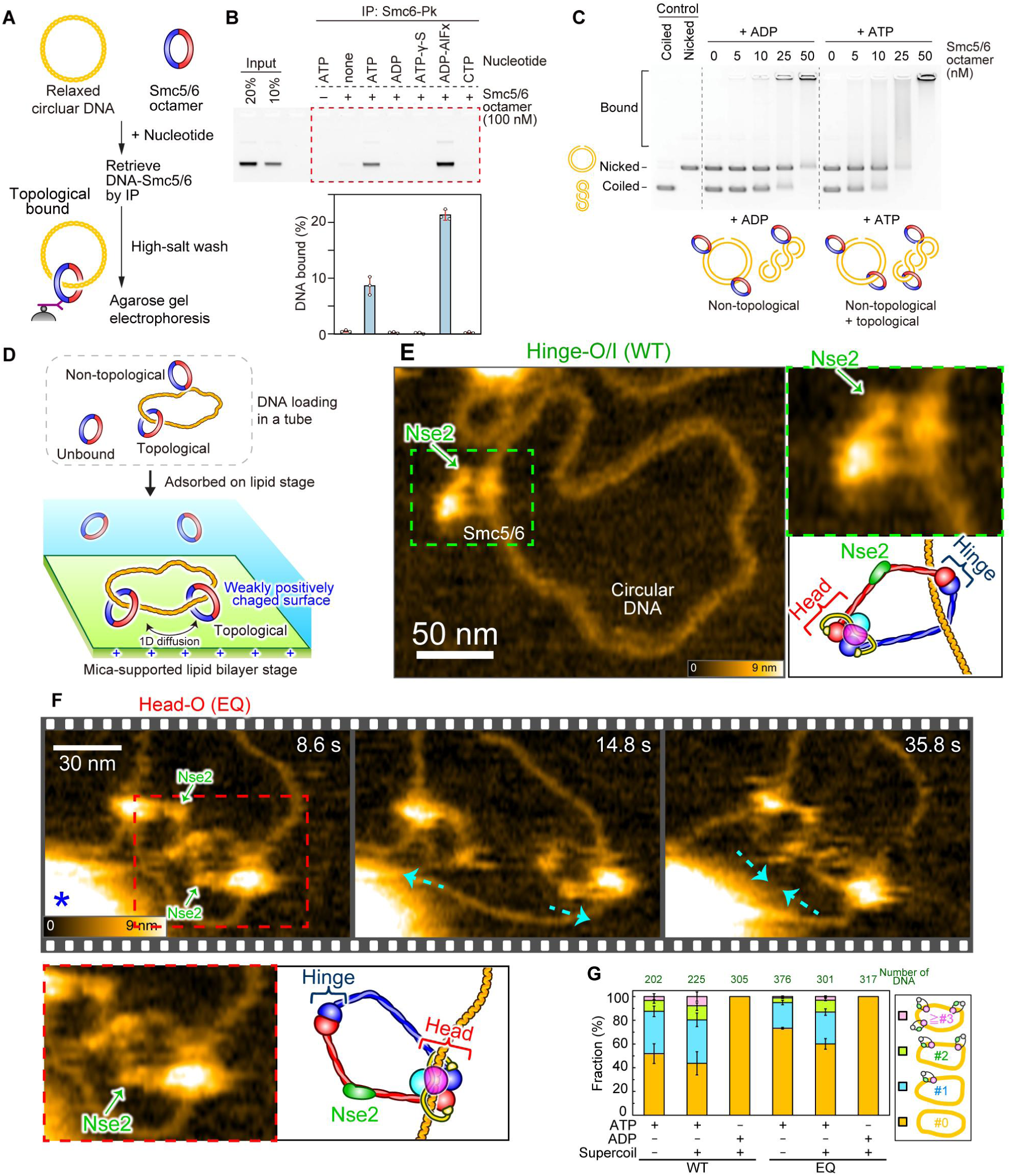
Visualization of DNA-bound Smc5/6 complex using the supported lipid bilayer stage. (**A**) Schematic of DNA loading assay. (**B**) Representative agarose gel analysis of the experiment as depicted in **a**. The graph presents the quantification of Smc5/6-bound DNA (obtained from three independent experiments; means ± s.d.). (**C**) ATP-independent DNA binding of Smc5/6 was analyzed using an electrophoretic mobility shift assay (EMSA). (**D**) Schematic of HS-AFM imaging using a positively charged lipid membrane as the imaging stage. (**E**) Overview scans of the DNA-bound wild-type Smc5/6 molecule, displaying the characteristic SMC ring structure (related to movie S4). Scan area, 350 × 280 nm^2^ at 200 × 80 pixels^2^, cropped to 260 × 206 nm². Imaging rate: 1 s per frame. (**F**) Successive HS-AFM images of the interaction between two EQ mutants bound to a DNA molecule. The cyan arrows indicate the direction of molecular movement as the molecules interact with each other, moving closer together and further apart. Scan area, 220 × 176 nm^2^ at 200 × 80 pixels^2^, cropped to 138 × 93 nm². Imaging rate: 0.6 s per frame. The blue asterisks indicate an aggregated structure of the lipid. (**G**) Quantification of Smc5/6 molecules bound to DNA (from three independent experiments), with error bars representing s.d..

To further assess DNA-binding activity, we performed an electrophoretic mobility shift assay using supercoiled and nicked (relaxed) circular DNA (Fig. 4C). This assay clearly indicated a distinct DNA shift in the presence of the Smc5/6 octamer and ATP. A weaker but still noticeable DNA shift was also observed in the presence of ADP, suggesting ATP-independent, non-topological DNA binding mediated by the DNA-binding motifs of the Smc5/6 complex (*46*). Additionally, Smc5/6 exhibited a relatively higher affinity for supercoiled DNA than for relaxed DNA. These results are consistent with previous biochemical studies (*20*, *24*).

Having established ATP-dependent Smc5/6 loading in bulk reactions, we next sought to visualize individual Smc5/6 molecules bound to DNA. After loading Smc5/6 octamer onto circular DNA (1895 bp, ∼644 nm) in the presence of ATP, the reaction mixture was applied onto the mica in a buffer containing potassium ions. However, under these conditions, multiple Smc5/6 molecules stacked on the DNA, as both DNA and Smc5/6 were adsorbed on the mica surface, making it difficult to track the dynamics of individual Smc5/6 on DNA (fig. S5C).

To overcome this difficulty, we adapted our previously developed supported lipid bilayer system, which features a weakly positively charged surface that adsorbs DNA while repelling proteins (Fig. 4D) (*37*, *38*). This approach successfully visualized both wild-type and EQ mutant Smc5/6 loaded onto DNA, revealing their characteristic ‘SMC’ shape along with Nse2 subdomain and freely dynamic behavior (Fig. 4E, F, fig. S6, movie S4). In most cases, a single Smc5/6 molecule bound to one DNA molecule, although in some instances, multiple Smc5/6 bound to the same DNA molecule (Fig. 4F, G, fig. S6, movie S5). We also observed that Smc5/6 bound to both supercoiled and relaxed DNA, with a slight preference for supercoiled DNA, consistent with the bulk biochemical experiments shown in Fig. 4C. In contrast to the presence of ATP, DNA-bound Smc5/6 was rarely detected in the presence of ADP (Fig. 4G). Although Smc5/6 exhibited ATP-independent, non-topological DNA binding in the bulk assay (Fig. 4C), this electrostatic DNA binding appeared to be weak and was therefore rarely observed under HS-AFM on the lipid stage. Given that topological Smc5/6 loading depends on ATP (Fig. 4B, fig. S5A, B) (*20*, *24*), the Smc5/6 molecules observed by HS-AFM likely entrapped DNA within the ring compartment. Consistent with this notion, Smc5/6 randomly slid along the DNA during scanning. Together, these results demonstrate that the current HS-AFM setup enables visualization of both the subdomain-level structure and the dynamics of Smc5/6 on DNA.

### Smc5/6 interacts with DNA through both the ATPase heads and hinge regions

Like other SMC complexes, the Smc5/6 ring, formed by Smc5, Smc6, and Nse4, is divided into two compartments, the SMC and the kleisin compartments, by the interaction of the ATPase heads (Fig. 5A). Accordingly, Smc5/6 should encircle DNA within either the SMC or kleisin compartment, or both (*24*). We observed two distinct types of DNA-bound Smc5/6 molecules. One type was associated with DNA at the hinge domain, exhibiting either the O-form or I-form structures (referred to hereafter as Hinge-O/I; Fig. 5B, movie S6). Approximately 50% of the wild-type Smc5/6 molecules were classified as Hinge-O/I (Fig. 5E). The other type bound to DNA at the ATPase heads, predominantly adopting the I-form structure (hereafter referred to as Head-I; Fig. 5C, movie S6). We noticed that, unlike the observations on mica (Fig. 2A), Nse2 was not detected in the DNA-bound, Head-I molecules. During real-time tracking, we occasionally observed transitions from the Head-I to the O-forms with the appearance of the Nse2 subunit (hereafter referred to as Head-O, fig. S7A, Fig. 5D). This suggests that DNA-bound Smc5/6 in the Head-I state was adsorbed onto the lipid membrane surface in a different orientation, probably with Nse2 facing down, compared to the observations on mica. Supporting this notion, Nse2 was not visible in the pseudo-AFM image when Nse2 was facing downwards in the apo-state atomic model (I-form) (fig. S7B, C).

**Figure 5.**
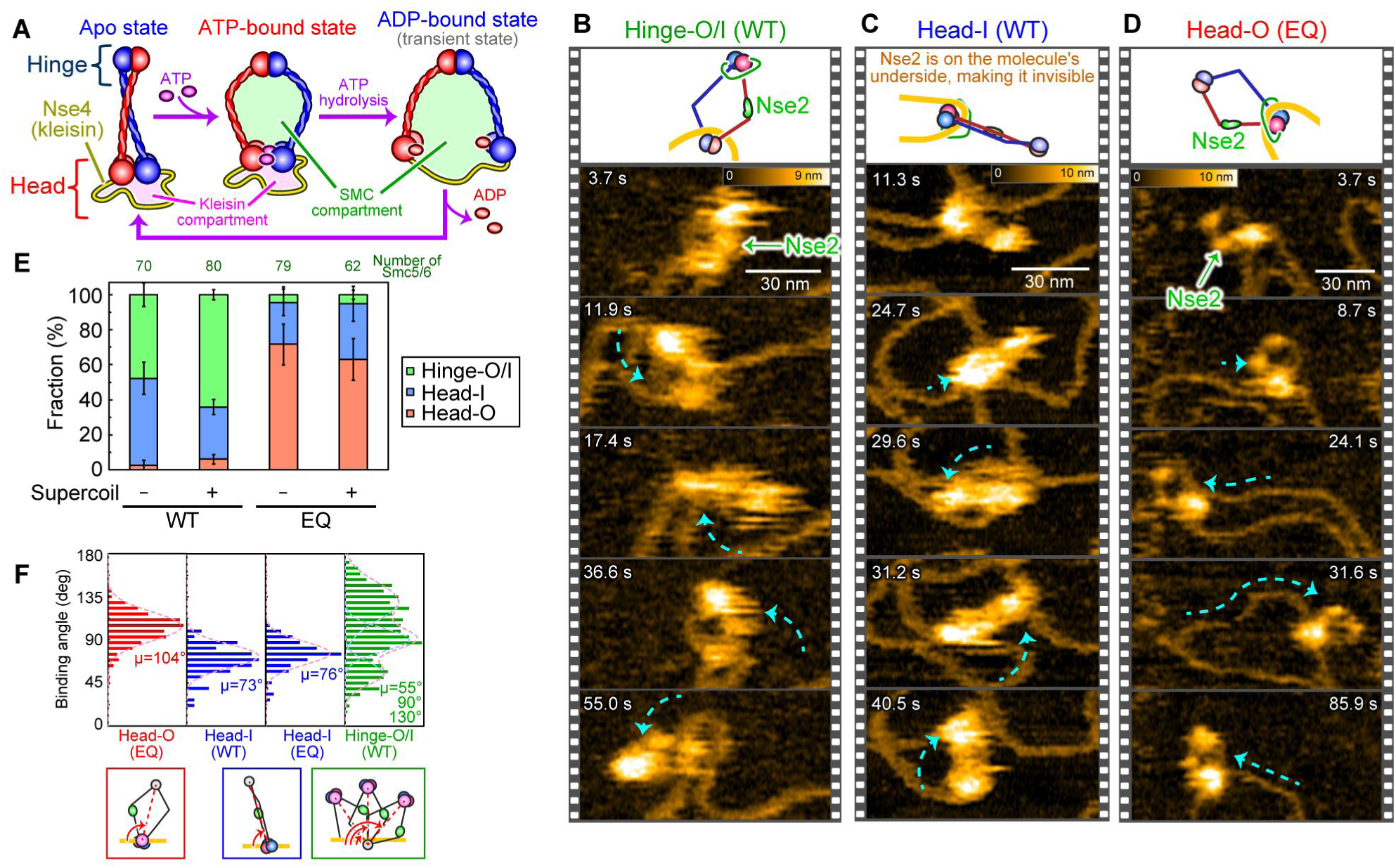
Classification of Smc5/6 molecules bound to DNA. (**A**) Schematics of the apo, ADP-bound, and ATP-bound Smc5/6 complex. The SMC ring is divided into the SMC and kleisin compartments. (**B**-**D**) Real-time imaging of three types of DNA-bound Smc5/6 molecules showing Hinge-O/I (B), Head-I (C), and Head-O (D) states (related to movie S6). The cyan arrows indicated the direction of the molecular movement. Scan area, 150 × 150 nm^2^ at 160 × 80 pixels^2^, cropped to 130 × 68 nm². f: Scan area, 150 × 120 nm^2^ at 160 × 64 pixels^2^, cropped to 128 × 71 nm². g: Scan area, 220 × 176 nm^2^ at 200 × 80 pixels^2^, cropped to 155 × 78 nm². Imaging rates: 0.37 s per frame (B), 0.34 s per frame (D), 0.6 s per frame (D). (**E**) Abundance ratios of DNA-bound Smc5/6 conformations (from three independent experiments). Error bars represent s.d.. (**F**) Distribution of Smc5/6 binding angles, analyzed from a total of 252, 55, 166, and 450 frames of successive HS-AFM images for EQ Head-O, WT Head-I, EQ Head-I, and WT Hinge-O/I, respectively (*n* = 3 molecules per condition).

In contrast to the wild-type, the majority (∼95%) of the EQ mutant bound to DNA at the ATPase heads, with the Hinge-O/I configuration being rarely observed. Furthermore, approximately 70% of the EQ mutant displayed the Head-O structure while bound to DNA (Fig. 5D, E, movie S6). These results suggest that the Head-O/I configurations sequester DNA within the kleisin compartment, whereas the Hinge-O/I configurations encircle DNA within the SMC compartment (see Discussion). Given that the EQ mutant, which stabilizes ATPase head engagement, primarily sequesters DNA at the ATPase head region, ATP hydrolysis appears to be essential for establishing the Hinge-O/I DNA-binding state.

During HS-AFM observation, the Smc5/6 complex in the Hinge-O/I configurations on DNA often exhibited swaying motion, whereas the head-bound molecules remained relatively stable. To quantify these differences in mobility, we measured the binding angles of Smc5/6 relative to DNA at each scanning time point (Fig. 5F, fig. S8A; see Materials and Methods). The histograms of the binding angles showed that the Head-I molecules were distributed in a single peak, bound at an average of 73° (76° for the EQ mutant). Similarly, the Head-O molecules derived from the EQ mutant showed a single peak distribution, but with a peak binding angle of 104°, suggesting that Head-O and Head-I are similarly tilted relative to 90° but in opposite directions. In contrast, Hinge-O/I from the wild-type complex exhibited a broad distribution of binding angles, which could be fitted to multiple Gaussian peaks at 55, 90, and 130°. These results suggest that Smc5/6 stably grasps the DNA strand at the ATPase head region, while DNA binding at the hinge domain is less rigid, allowing Hinge-O/I molecules to exhibit greater flexibility. Additionally, we measured the angles of the DNA curvature at Smc5/6 binding sites (fig. S8B, C). The binding sites of Hinge-O/I molecules exhibited larger curvature angles (median at 110°) compared to those of Head-O/I molecules (less than 50°). These differences indicate that the Hinge-O/I preferentially associates with curved regions of DNA, whereas the head-binding Smc5/6 molecules bind to less curved regions, reflecting distinct DNA binding states.

### Smc5/6 modulates DNA structure

Single-molecule fluorescence imaging and biophysical studies have demonstrated that Smc5/6 compacts DNA strands in an ATP-dependent manner (*7*, *20*, *21*). As shown in Fig. 6A, we noticed that the nicked circular DNA substrate, which typically appears as an extended circle strand under HS-AFM (–Smc5/6, Fig. 6A–a1), tended to become highly deformed upon binding of Smc5/6. We classified the Smc5/6-bound DNA molecules into the following three types: Smc5/6 bound to one DNA segment (Bound 1, Fig. 6A–a2), Smc5/6 interacting with two DNA segments (Bound 2, Fig. 6A–a3), and Smc5/6 binding at twisted regions of the DNA (Twisted, Fig. 6A–a4). For the “Bound” species, we primarily observed “Bound 1” (15 out of 17 molecules), with fewer Smc5/6-bound DNA molecules classified as “Bound 2”. Subsequent real-time tracking revealed that the “Bound 2” state was transient, with Smc5/6 tending to bridge then release two segments of DNA (fig. S9).

**Figure 6.**
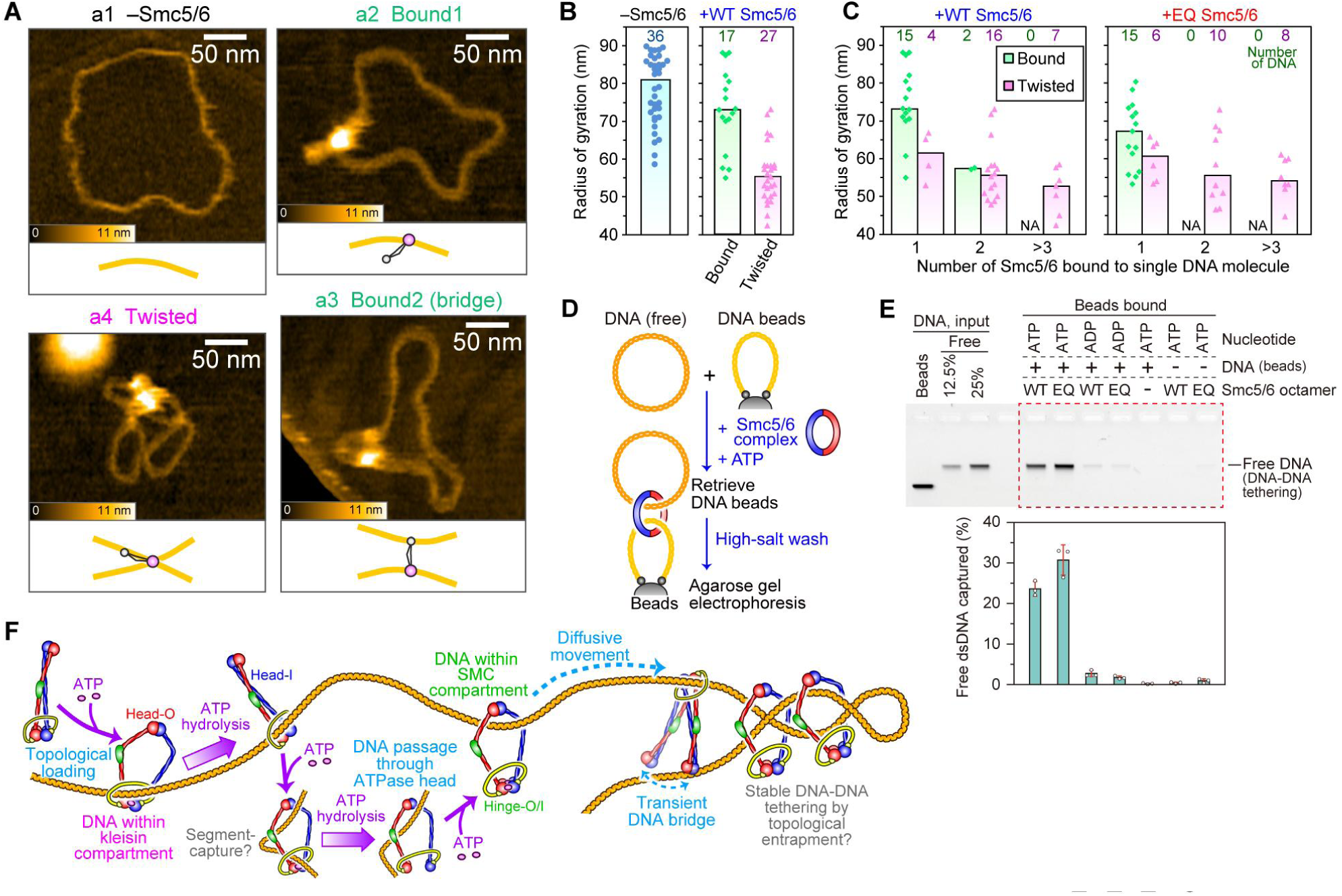
DNA structures modulated by Smc5/6 tethering activity. (**A**) Representative HS-AFM images of Smc5/6 bound to nicked, relaxed circular DNA substrates on a lipid membrane. The four binding conformations are shown: –Smc5/6 (a1), bound1 (a2), bound2 (tethered) (a3), and twisted (a4). a1: Scan area, 350 × 280 nm^2^ at 200 × 80 pixels^2^, cropped to 280 × 252 nm². a2: Scan area, 450 × 360 nm^2^ at 240 × 96 pixels^2^, cropped to 315 × 245 nm². a3: Scan area, 350 × 280 nm^2^ at 200 × 80 pixels^2^, cropped to 301 × 252 nm². a4: Scan area, 350 × 280 nm^2^ at 200 × 80 pixels^2^, cropped to 283 × 224 nm². Imaging rates: 1 s per frame (a1, a4), 1.5 s per frame (a2), 0.9 s per frame (a3). (**B**) Quantification of the radius of gyration of Smc5/6-bound DNA. (**C**) Same as **b**, except that Smc5/6-bound DNAs were further classified according to the number of Smc5/6 molecules. (**D, E**) Schematic (D) and representative agarose gel image (E) of the DNA–DNA tethering assay. The graph shows the quantification of captured free DNA (from three independent experiments; means ± s.d.). (**F**) A schematic model depicting the ATP-dependent transactions of Smc5/6 on DNA, including topological loading, diffusive movement and DNA-DNA tethering.

In the “Twisted” DNA species, Smc5/6 appeared to interact with more than one DNA segment. To quantify the degree of compaction, we calculated the radius of gyration (RoG) of the “Bound” and “Twisted” DNA molecules. As shown in Fig. 6B, the mean RoGs of both the “Bound” (∼75 nm) and “Twisted” (∼55 nm) species were smaller than that of the naked DNA substrate (∼80 nm), with a higher degree of DNA compaction for the “Twisted” species. Interestingly, we also found that multiple Smc5/6 molecules tended to bind to the “Twisted” species more frequently than the “Bound” species (Fig. 6C). These results suggest that Smc5/6 mediates DNA compaction through DNA twisting, particularly when multiple Smc5/6 molecules are loaded onto DNA. Notably, similar DNA compaction was observed with the EQ mutant (Fig. 6C), indicating that this compaction does not rely on ATP hydrolysis. This property contrasts with DNA loop extrusion, which has been reported to require continuous ATP hydrolysis by SMC complexes (*4–7*).

As Smc5/6 was frequently observed in contact with two DNA strands during our HS-AFM analyses, we speculated that DNA compaction might be facilitated by tethering two independent DNA strands. In addition to loop extrusion, cohesin and condensin have also been shown to tether two separate DNA strands in an ATP-dependent manner (*9–11*). To test whether Smc5/6 mediates DNA–DNA tethering, a linear DNA fragment biotinylated at both ends was immobilized on streptavidin-conjugated magnetic beads (Fig. 6D). These DNA-coated beads were then incubated with circular plasmid DNA (free DNA) and Smc5/6 in the presence of ATP. After a high-salt wash, the captured free circular DNA was analyzed by agarose gel electrophoresis. Approximately 25% of the input free DNA was recovered in the presence of ATP (Fig. 6E, WT). In contrast, recovery of free DNA was less than 3% when ATP was replaced by ADP or omitted. These results indicate that Smc5/6 mediates DNA–DNA tethering in an ATP-dependent manner. Efficient recoveries of free DNA were also seen using the EQ mutant with ATP (Fig. 6E, EQ) or WT with ADP-AlF_x_ (fig. S10A), indicating that the observed DNA–DNA tethering does not require ATP hydrolysis.

To investigate whether the observed DNA tethering was mediated by topological entrapment, we digested free DNA at one site using a restriction enzyme after recovery of the DNA tethering products (fig. S10B). This released linearized free DNA into the supernatant (fig. S10C, D). Similarly, single digestion of bead-attached DNA released intact free DNA from the beads and simultaneously dissociated Smc5/6 from the DNA (fig. S10C, D). These bulk biochemical experiments suggest that Smc5/6 mediates mediate DNA–DNA tethering via topological DNA entrapment.

## Discussion

In this study, we visualized the structural dynamics of budding yeast Smc5/6 complexes at submolecular resolution using HS-AFM under a physiological buffer condition. Using the Smc5/6 hexamer, we primarily observed two types of extended molecules, the I- and O-forms, depending on their ATP binding state. These structural variations are distinct from those of cohesin and condensin, which display sharply folded conformations by bending the SMC arms at their elbows in a manner not seen with Smc5/6 (*32*, *33*, *47–51*). Our data show that Nse2 maintains the extended conformation of Smc5/6 by preventing bending at the elbows of its SMC arms, consistent with recent electron microscopic observations of Smc5/6 lacking Nse2 (*28*). As Nse2 is essential for cell viability (*52*), the observed structural constraint of Smc5/6 may be related to its chromosomal functions, probably through its DNA-binding strategies observed in this study.

By optimizing the experimental conditions, namely through the use of the supported lipid bilayer system (*37*, *38*) which minimizes the surface adsorption of Smc5/6, we were able to visualize the molecular dynamics of Smc5/6 bound to DNA with unprecedented spatiotemporal resolution. This result revealed the existence of three distinct types of DNA-bound Smc5/6. Both the Head-O (EQ mutant) and Head-I (wild-type) molecules bind to DNA at the ATPase head region, whereas the Hinge-O/I molecules, observed mainly with wild-type Smc5/6, hold DNA within the SMC compartment.

We speculate that Smc5/6 initially entraps DNA within the kleisin compartment and subsequently transfers it to the SMC compartment upon ATP hydrolysis (Fig 6F, fig. S11). The Head-O molecule, frequently observed with the EQ mutant, represents the state immediately after Smc5/6 topologically entraps DNA (Fig. 6F, Topological loading). A recent biochemical study using protein crosslinking experiments has shown that Smc5/6 topologically entraps DNA through transient opening of the Smc6–Nse4 interface (N-gate) (*24*). In the non-ATP binding apo state, Nse5–Nse6 mediates the N-gate opening by inhibiting the interaction between Nse4 and Smc6 (*24*, *27*, *28*). The reclosure of the N-gate is facilitated by the ATP-dependent engagement of the SMC ATPase heads and DNA binding, leading to the topological capture of DNA within the kleisin compartment (*24*, *28*). Subsequent ATP hydrolysis destabilizes and converts the O-form to the I-form, which is commonly observed with the wild-type complex (Fig 6F, Head-I).

How does Smc5/6 ultimately deliver DNA to the SMC compartment? A recent cryo-EM study of fission yeast cohesin suggests that it initially entraps DNA via the N-gate by forming the DNA-gripping intermediate (*49*). In this state, DNA is also positioned at the top of the ATPase heads, along with the cohesin loader. A similar intermediate, termed the segment-capture state, has been proposed for Smc5/6, in which a small DNA loop is topologically entrapped within the Nse4 kleisin compartment, while being held by the SMC compartment (*24*) (Fig. 6F, Segment-capture). We speculate that the segment-capture state is relatively unstable and predisposed to transition into the Hinge-O/I or Head-O/I state. ATP hydrolysis induces disengagement of the ATPase heads, allowing the DNA segment located above the head to pass through, thereby forming the Hinge-O/I state (Fig. 6F, DNA within the SMC compartment). Conversely, slippage of a small DNA loop from the SMC compartment without the head disengagement leads to the formation of the Head-O/I states. Recent cryo-EM studies of the prokaryotic SMC complexes have revealed the DNA-holding state, where DNA is trapped within the kleisin compartment but not within the SMC compartment (*53*, *54*). The Head-O/I Smc5/6 molecules may represent a similar DNA-holding mode. Once DNA is entrapped, Smc5/6 exhibits diffusive motion along the DNA, a property that may facilitate its translocation along chromatin from its initial loading site (Fig. 6F, Diffusive movement).

In addition to topological DNA entrapment, Smc5/6 has been shown to form DNA loop through extrusion (*7*). Cohesin and condensin have been proposed to mediate DNA loop extrusion via bending and stretching motions of their SMC arms at the elbows (*32*, *33*, *55*, *56*). In contrast, Nse2, which restricts the Smc5/6 arm bending in our HS-AFM observations, has paradoxically been reported to essential for Smc5/6-mediated loop extrusion (*7*). In addition, Smc5/6 facilitates loop extrusion via dimer formation (*7*). These observations suggest that Smc5/6 may employ a distinct mechanism of DNA loop extrusion compared to cohesin and condensin. Whether and how the Nse2-regulated conformation of Smc5/6 is linked to both DNA loop extrusion and topological DNA entrapment remains an important topic for further investigation.

Our HS-AFM observations also revealed that Smc5/6-bound DNA molecules, especially those with multiple Smc5/6 molecules, tended to be twisted, leading to DNA compaction. One possible mechanism for this observed DNA compaction is DNA loop extrusion by Smc5/6, which has been reported to require continuous ATP hydrolysis (*7*). However, we observed a similar level of DNA compaction with the EQ mutant, suggesting that efficient ATP hydrolysis is not required for the compaction observed in this study. Furthermore, our biochemical experiments also detected physical tethering of two independent DNA molecules in an ATP-binding-dependent, but not hydrolysis-dependent manner. Based on these findings, we speculate that Smc5/6 compacts DNA by a DNA–DNA tethering mechanism (Fig. 6F, DNA–DNA tethering).

Previous force measurement studies using magnetic tweezers have shown that Smc5/6 mediates more efficient DNA compaction when two DNA strands are aligned in close proximity (*20*). Thus, Smc5/6 may associate more stably with DNA in the tethered state. Interestingly, cohesin and condensin in their monomeric form have been shown to capture a second DNA molecule by topological entrapment (*9–11*), suggesting that DNA–DNA tethering activity is a conserved biochemical property among SMC complexes, in addition to DNA loop extrusion. The mode by which Smc5/6 molecules associate with DNA at the twisted region remains unclear due to the limited structural resolution in HS-AFM. In the future, it will be of great interest to investigate whether, and how, Smc5/6 employs the DNA-binding strategies we observed to recognize and manipulate higher-order and intrinsically harmful DNA structures that arise during the processes of DNA repair and transcription.

## Acknowledgements

We thank Prof. Toshio Ando (Kanazawa University) and Prof. Takayuki Uchihashi (Nagoya University) for their technical assistance, Dr. Hirotatsu Imai (University of the Ryukyus) for discussion on biological experiments, Dr. Steven J. McArthur (Kanazawa University) and Dr. Frank Uhlmann (The Francis Crick Institute) for their critical reading of the manuscript.

## Funding

PRESTO, Japan Science and Technology Agency (JST), JPMJPR20E3 and JPMJPR23J2 (KU).

PRESTO, JST, JPMJPR19KB (YM)

CREST, JST, JPMJCR1762 (NK)

KAKENHI, Japan Society for the Promotion of Science (JSPS), 19K15409 and 21K04849 (KU)

KAKENHI, JSPS, 26119003 and 20H00327 and 24H00402 (NK)

KAKENHI, JSPS, 23K23814 and 24H02282 (YM);

The Takeda Science Foundation (YM)

The World Premier International Research Center Initiative (WPI), MEXT (NK)

Author contributions

Conceptualization: KU, YM

Methodology: KU, YK, YM, NK

Investigation: KU, YK, YM

Visualization: KU, YK, YM

Funding acquisition: KU, YM, NK

Project administration: YM, NK

Supervision: YM, NK

Writing – original draft: KU, YM

Writing – review & editing: KU, YK, YM, NK

## Competing interests

The authors declare no competing interests.

## Data and material availability

The data that support the findings of this study are available from the corresponding author upon reasonable request (Kenichi Umeda), Yasuto Murayama and Noriyuki Kodera.

## Supplementary Materials

Materials and Methods

Figs. S1 to S11

Table S1 to S3

Movie S1 to S6

References (57–60)

## Notes

### Competing Interest Statement

The authors have declared no competing interest.

