## Supplementary Materials for "Submolecular video-imaging of the Smc5/6 complex topologically bound to DNA"

**The PDF file includes:**

Materials and Methods
Figs. S1 to S11
Tables S1 to S3
References (57–60)

**Other Supplementary Materials for this manuscript include the following:**

Movies S1 to S6

### Materials and Methods

#### Cell culture media

Culture media used for protein expressions are listed below:

YP-raffinose medium: 1% yeast extract, 2% peptone, 1.5% raffinose, supplemented with 100 μg $mL^{-1}$ adenine

Terrific-broth medium: 1.2% tryptone, 2.4% yeast extract, 0.4% glycerol, 17 mM $NH_2PO_4$ and 72 mM $K_2HPO_4$

#### Protein Purification Buffers

Buffer used for protein purifications are listed below:

Cell lysis buffer: 40 mM Tris-HCl (pH 7.5), 0.5 mM tris(2-carboxyethyl)phosphine (TCEP), 10% (w/v) glycerol, 0.3 M NaCl

R buffer: 20 mM Tris-HCl (pH 7.5), 0.5 mM TCEP, 10% (w/v) glycerol

R1 buffer: 20 mM Tris-HCl (pH 7.5), 0.5 mM TCEP, 10% (w/v) glycerol, 0.4 M NaCl, 0.005% (w/v) Tween 20

R2 buffer: 20 mM Tris-HCl (pH 7.5), 0.5 mM TCEP, 10% (w/v) glycerol, 0.4 M NaCl

R3 buffer: 20 mM Tris-HCl (pH 7.5), 0.5 mM TCEP, 10% (w/v) glycerol, 0.4 M NaCl and 0.01% (w/v) Tween 20

R4 buffer: 20 mM Tris-HCl (pH 7.5), 0.5 mM TCEP, 10% (w/v) glycerol, 0.3 M NaCl, 0.005% (w/v) Tween 20

#### Purification of the budding yeast Smc5/6 complex

Complementary DNAs (cDNA) encoding the budding yeast Smc5, Smc6, Nse1, Nse2, Nse3, Nse4, Nse5, and Nse6 were amplified by PCR from *Saccharomyces cerevisiae* W303. The resultant cDNAs were cloned into yeast shuttle vectors under the control of the *GAL1* or *GAL10* promoter. The Smc6-encoding cDNA, fused to 3× Pk epitope tag and a protein A tag at the C-terminus, separated by a PreScission protease recognition sequence, along with the *SMC5* cDNA, was cloned into the YIplac211 (*URA3*) vector. The cDNA encoding Nse1, fused to a 7× histidine tag at the N-terminus, and the *NSE3* cDNA were cloned into the YIplac128 (*LEU2*) vector. The Nse4-encoding cDNA, fused to a 7× histidine tag at the C-terminus, and the *NSE2* cDNA were cloned into the YIplac204 (*TRP1*) vector. The Nse6-encoding cDNA, fused to a 7× histidine tag at the N-terminus, and the *NSE5* cDNA were cloned into the pRSII402 (*ADE2*) vector. The linearized DNA constructs were sequentially integrated into the budding yeast strain (*MATa, ade2-1, trp1-1, can1-100, leu2,3,112, his3-11,15, ura3-52, pep4Δ::HIS3MX*) at the *URA3*, *LEU2*, *TRP1* and *ADE2* loci, respectively.

The resultant Smc5/6 octamer (comprising Smc5, Smc6, Nse1, Nse2, Nse3, Nse4, Nse5, and Nse6) expression yeast was grown in 7.5 L (1.5 L × 5 cultures) of YP-raffinose media at 30 ºC to a cell density of 1–3 × $10^7$ per mL. Galactose was then added to the cell culture at a final concentration of 2% and the cells were grown further at 30 ºC for 2 h to induce protein expression. The cells were collected by centrifugation and washed once with ice-chilled cell lysis buffer and then resuspended in one quarter volume of the same buffer containing 0.5 mM phenylmethylsulphonyl fluoride (PMSF). The cell suspension was dropped directly into liquid nitrogen for freezing and then ground in a freezer mill (SPEX CertiPrep 6850).

All subsequent procedures were carried out at 4 ºC. The resultant cell powder was thawed at 4 ºC and resuspended for 30 min in 3 volumes of cell lysis buffer containing 0.005% (w/v) Tween 20, 0.5 mM PMSF and cOmplete protease inhibitor cocktail (Merck). After clarification by centrifugation at 28,000 × *g* for 45 min, the cell lysate was loaded onto 1.5 mL of human IgG agarose (Sigma) at 0.75 mL $min^{-1}$ using a peristaltic pump. The beads were washed with 20 mL of R1 buffer containing 0.5 mM PMSF, then 80 mL of R2 buffer. The beads were resuspended in the same buffer with 1 mM TCEP, 10 µg $mL^{-1}$ RNase A and 5 U $mL^{-1}$ of PreScission protease and incubated overnight. The eluate was passed through 0.2 mL of glutathione Sepharose (Cytiva) to remove the PreScission protease. This fraction was then diluted with R buffer to a final concentration of 0.2 M NaCl and applied to a heparin HP column (1 mL, Cytiva). The column was developed with a linear gradient of 0.2 to 1 M KCl in R buffer. Peak fractions (~0.35 M KCl) were pooled and concentrated by ultrafiltration (Amicon Ultra, 30K, Millipore), then loaded onto a Superose 6 Increase 10/300 GL gel filtration column (24 mL, Cytiva), which was equilibrated in R buffer containing 0.3 M KCl. Peak fractions were concentrated by ultrafiltration, and the aliquots were snap-frozen in liquid nitrogen and stored at –80 ºC. Smc5/6 hexamers (Smc5, Smc6, Nse1, Nse2, Nse3, and Nse4) and the EQ mutants (Smc5 E1015Q, Smc6 E1048Q) were expressed and purified using the same procedure.

**Purification of the budding yeast Nse5–Nse6 heterodimer**

The Nse5–Nse6 complex was expressed and purified in *Escherichia coli*. as described previously (*41*) with the following modification. Complementary DNAs (cDNA) encoding the budding yeast Nse5 and Nse6 were amplified by PCR from *Saccharomyces cerevisiae* W303. The Nse5-encoding cDNA fused to a PreScission protease recognition sequence and a 2× Strep tag at the C-terminus and the *NSE6* cDNA, both of which were connected with the ribosome binding sequence, were cloned into the *E. coli* protein expression vector pET21a under the control of a T7 promoter. The resultant Nse5–Nse6 expression plasmid was introduced into the *E. coli* Rosetta 2 (DE3). The transformants were grown in 3 L (1.5 L ×2 cultures) of Terrific-broth medium containing 100 µg $mL^{-1}$ ampicillin and 17 µg $mL^{-1}$ chloramphenicol at 37 ºC to an optical density of 0.6 at 600 nm. Cell cultures were cooled to 20 ºC and further cultivated for 16 h after the addition of isopropyl β-D-1-thiogalactopyranoside (IPTG) to a final concentration of 0.25 mM to induce protein expression.

All subsequent procedures were carried out at 4 ºC. Cells were collected by centrifugation and washed once with ice-chilled R2 buffer. The cell pellets were resuspended in 5 volumes of the same buffer supplemented with 1 mM PMSF and 1 mM 4-(2-aminoethyl)benzenesulphonyl fluoride hydrochloride (AEBSF). After the addition of one tenth volume of 10× BugBuster solution (Merck), the cells were disrupted by sonication. After clarification by centrifugation at 28,000 × *g* for 45 min, the cell lysate was passed through a 0.22 µm filter and then loaded onto 2 mL of Strep-Tactin agarose (IBA) at 0.75 mL $min^{-1}$ using a peristaltic pump.

The beads were washed sequentially with 40 mL of R3 buffer, 20 mL of R4 buffer containing 1 mM ATP and 5 mM $MgCl_2$ and then 60 mL of R4 buffer. The beads were further washed with 5 mL of R4 buffer containing 2.5 mM TCEP and then resuspended in 1.5 mL of the same buffer containing 5 U $mL^{-1}$ of PreScission protease and incubated overnight. The eluate was passed through 0.2 mL of glutathione-Sepharose (Cytiva) to remove the PreScission protease. This fraction was then diluted with R buffer to a final NaCl concentration of 0.15 M and applied to a heparin HP column (1 mL, Cytiva), which was developed with a linear gradient of 0.1–1 M NaCl in R buffer. Peak fractions (eluting around ~0.3 M NaCl) were pooled and concentrated by ultrafiltration (Amicon Ultra, 30K, Millipore), then applied to a Superdex 200 Increase 10/300 GL column (24 mL, Cytiva), equilibrated in R buffer containing 0.2 M NaCl. Peak fractions were again concentrated by ultrafiltration, and aliquots were snap-frozen in liquid nitrogen and stored at –80 ºC.

**DNA substrates**

The 1895 bp plasmid DNA, pYST400, used for HS-AFM analysis was generated by ligation of the annealed oligonucleotides (5′-TATGCTGAGGTACGATATCTTA-3′ and its complementary oligo DNA) with the digested DNA fragment of pBluescript KSII (+) with *Ssp*I and *Pvu*II. Covalently closed circular plasmid DNA (cccDNA) of pYST400 (1895 bp), pBluescript KSII (+) (3.0 kbp) and pSKsxAS (4.3 kbp) (*9*) and pKSII-E2 (7.8 kbp) (*9*) was prepared by the plasmid purification kit (Maxi prep. Kit, Qiagen). Relaxed circular DNA (rcDNA) was prepared by treating cccDNA with *E. coli* topoisomerase I (NEB). Nicked circular DNA (ncDNA) of pYST400 or pBluescript KSII (+) was prepared using *Nt.Bbv*CI or *Nb.Bss*SI, respectively. To prepare immobilized, closed topology DNA, the 3.0 kbp linear DNA was first prepared by PCR amplification with a pair of 5′ biotinylated DNA primers (759 and 760 bp) using pKSII-E2 as the template. The resultant linear DNA was mixed and immobilized on streptavidin conjugated magnetic beads (Thermo), as described in Ref 9.

**Buffers used for the biochemical and HS-AFM experiments.**

Buffer used for the biochemical and HS-AFM experiments in this study are listed below:

| | |
|---|---|
| ATPase reaction buffer: | 25 mM HEPES-KOH (pH 7.5), 1 mM TCEP, 5% (w/v) glycerol, 75 mM potassium acetate (KOAc), 30 mM KCl, 1 mM magnesium acetate ($Mg(OAc)_2$), 0.01% (w/v) NP-40 |
| SL buffer: | 25 mM HEPES-KOH (pH 7.5), 1 mM TCEP, 5% (w/v) glycerol, 75 mM KOAc, 30 mM KCl, 1 mM $Mg(OAc)_2$, 0.05% (w/v) NP-40 |
| IP500 buffer: | 25 mM HEPES-KOH (pH 7.5), 500 mM NaCl, 0.25% (w/v) NP-40, 1 mM ethylenediaminetetraacetic acid (EDTA) |
| IP100 buffer: | 25 mM HEPES-KOH (pH 7.5), 100 mM NaCl and 0.1% (w/v) NP-40 |
| Elution buffer: | 10 mM Tris-HCl (pH 7.5), 0.5% (w/v) sodium dodecyl sulfate (SDS), 1 mM EDTA, 1 mg $mL^{-1}$ proteinase K |

| | |
|---|---|
| RE buffer: | 25 mM Tris-HCl (pH 7.5), 1 mM TCEP, 10 mM $MgCl_2$, 100 mM NaCl and 0.1% (w/v) NP-40 |
| DC500 buffer: | 25 mM HEPES-KOH (pH 7.5), 500 mM NaCl, 1 mM EDTA, 0.05% (w/v) NP-40 |
| DC100 buffer: | 25 mM HEPES-KOH (pH 7.5), 100 mM NaCl, 0.05% (w/v) NP-40 |
| AFM-A buffer: | 25 mM HEPES-KOH (pH 7.5), 1 mM TCEP, 10% (w/v) glycerol, 300 mM KOAc, 1 mM $Mg(OAc)_2$ |
| AFM-B buffer: | 25 mM HEPES-KOH (pH 7.5), 1 mM TCEP, 5% (w/v) glycerol, 75 mM KOAc, 1 mM $Mg(OAc)_2$ |
| AFM-C buffer: | 25 mM HEPES-KOH (pH 7.5), 1 mM TCEP, 10% (w/v) glycerol, 30 mM KOAc, 1 mM $Mg(OAc)_2$ |

**ATPase assay**

Purified budding yeast Smc5/6 (100 nM) was mixed with pBluescript KSII rcDNA (16.6 nM) on ice in 20 µL of ATPase reaction buffer. The reaction was initiated by adding ATP (0.5 mM) and incubated at 30 ºC. 4 µL of samples were then taken at each time-point and placed on ice to stop the reaction. Generation of inorganic phosphate on ATP hydrolysis was measured using a commercially available phosphate-detection kit (BioAssay Systems).

**Electrophoretic mobility shift assay**

Smc5/6 was mixed with both cccDNA and ncDNA from pBluescript KSII (0.8 nM) at increasing concentrations in 15 µL of SL buffer, containing 5 mM ATP or ADP, and incubated at 30 ºC for 15 min. Samples were analyzed by 0.8% agarose gel electrophoresis in TAE (pH 7.5) buffer at room temperature (~23 ºC) for 90 min at 3.6 V $cm^{-1}$, followed by Sybr Gold staining. Gel images were captured using an Image Quant LAS-400 mini gel documentation system (Fujifilm).

**Smc5/6 loading assay**

Smc5/6 was mixed on ice in 15 µL of SL buffer, either in the absence or presence of 5 mM ATP, ADP, CTP, ATP-γ-S or 5 mM ADP supplemented with 1 mM $AlCl_3$ and 10 mM NaF. The reaction was initiated by the addition of ncDNA of pBluescript KSII (3.3 nM) and incubated at 30 ºC for 30 min. The reaction was terminated by adding ice-chilled IP500 buffer. Anti-Pk (V5, Bio-Rad), bound to protein A-conjugated magnetic beads (ThermoFisher), was added to the reaction mixture and rocked at 4 ºC overnight (~15 h). The beads were washed three times with 750 µL of IP500 buffer and then once with IP100 buffer. The washed beads were resuspended in elution buffer and incubated at 37 ºC for 20 min. The recovered DNA was separated by 1% agarose gel electrophoresis in TAE buffer (pH 8.2) at room temperature (~23 ºC) for 60 min at 3.6 V $cm^{-1}$, followed by Sybr Gold staining. Gel images were captured using an Image Quant LAS-400 mini gel documentation system (Fujifilm), and band intensities were quantified using MultiGauge software (Fujifilm).

In the experiment involving linearization of Smc5/6-bound DNA, the loading reaction and subsequent immunoprecipitation were carried out as described above using cccDNA of pBluescript KSII (+). The resultant beads were further washed with RE buffer. The washed beads were incubated with *Pst*I (20 U, Takara) in 10 μL of RE buffer at 7 ºC for 20 min. The NaCl concentration was adjusted to 500 mM in 15 μL, and incubated for an additional 15 min on ice. DNA in both the supernatant and bead fractions was analyzed as described above.

**DNA–DNA tethering assay**

The concentrations indicated are final concentrations in the total reaction volume (15 μL). Smc5/6 (100 nM) was initially mixed on ice in 10 μL of SL buffer containing the indicated adenosine derivatives (5 mM). The reaction was initiated by adding 5 μL of a mixture of rcDNA (pSKsxAS, 4.3 kbp, 1.67 nM) and the immobilized, closed-topology DNA beads (3.0 kbp, 1.16 nM) in SL buffer. The mixture was incubated at 30 ºC for 30 min with occasional agitation. The beads were then resuspended in 150 μL of DC500 buffer and further incubated at 30 ºC for 3 min with occasional agitation. After washing the beads once with DC500 buffer and then once with DC100 buffer, the recovered beads were suspended in 15 μL of elution solution and incubated at 37 ºC for 20 min. The recovered DNA was separated by 1% agarose gel electrophoresis in TAE buffer (pH 8.2) at room temperature (~23 ºC) for 70 min at 3.6 V $cm^{-1}$, and stained with Sybr Gold.

In the experiment including linearization of captured DNA by Smc5/6, the DNA tethering reaction and subsequent high-salt wash were performed as described above. The resultant beads were further washed with RE buffer, then resuspended and further incubated at 7 ºC for 20 min in 10 μL of the same buffer containing *Pst*I (20 U, Takara) to digest rcDNA (pSKsxAS) or *Pfl*MI (20 U, NEB) to digest immobilized DNA at a single locus, respectively. The NaCl concentration was adjusted to 500 mM in 15 μL and incubated for an additional 15 min on ice. DNA in the supernatant and beads fractions was analyzed as described above, using elution solution without proteinase K. Smc5/6 in both fractions were also analyzed by SDS-PAGE followed by western blotting using an anti-Pk antibody to detect Smc6.

**High-speed AFM setup**

As described in our previous studies (*38*), we employed a lab-built high-speed AFM equipped with a commercially available cantilever specialized for bio-imaging (Olympus: BL-AC10DS-A2). The resonance frequency, spring constant, and the quality factor of the cantilever in a buffer solution were approximately 500 kHz, 0.1 N $m^{-1}$, and 1.5, respectively. An amorphous carbon tip, approximately 500 nm in length, was fabricated on the cantilever bird's beak by electron beam deposition. The free oscillation amplitude of the cantilever was a few $nm_{p\text{-}0}$, with the setpoint amplitude set to around 90% of the free amplitude.

In our previous setup, we have employed full-cycle digital lock-in detection using a field-programmable gate array to measure the cantilever amplitude. To increase the detection bandwidth, we implemented a half-cycle detector (*57*), which enabled imaging without damaging molecules even during high-speed scanning. We used control software developed in Visual Studio for HS-AFM control (*58*). To quantitatively measure the molecular size and DNA length, we implemented a function to compensate for the nonlinearity and tip-position dependence of the XY piezo scanner (this will be published elsewhere).

**High-speed AFM imaging**

For observations of Smc5/6 using a mica imaging stage, proteins were incubated in AFM-A buffer in the absence or presence of 5 mM ATP or ADP on ice for 5 min. Then, 1 μL of the protein mixture was deposited on a freshly cleaved mica surface and incubated for 5 min before washing away unbound proteins with the same buffer. HS-AFM observations were performed in the same solution. Although approximately 80–90% of the observed molecules were imaged in an intact state, the remaining molecules showed dissociation of either the head or hinge domains at the start of observation or within the first few frames. Only molecules that remained intact throughout imaging for several tens of frames were included in the analysis shown in Figs. 1 and 3.

To observe DNA-bound Smc5/6, the Smc5/6 octamer (50 nM) was first incubated with 1895 bp circular DNA (5.3 nM, pYST400) in AFM-B buffer containing 5 mM ATP at 30 °C for 2 h. For HS-AFM observation of DNA-bound Smc5/6 using a mica imaging stage, 1 μL of the DNA loading reaction was deposited on a freshly cleaved mica surface and incubated for 5 min. After rinsing with AFM-C buffer containing 5 mM ATP, HS-AFM observation was performed in the same buffer condition.

For observations of DNA-bound Smc5/6 using a mica-supported lipid bilayer imaging stage, 1,2-dipalmitoyl-*sn*-glycero-3-phosphocholine (DPPC; Avanti Polar Lipids) and 1,2-dipalmitoyl-3-trimethylammonium-propane (DPTAP; Avanti Polar Lipids), dissolved in chloroform, were mixed at a weight ratio of 9:1. The chloroform was then evaporated to form a lipid film, which was subsequently hydrated in water to prepare liposomes. This preparation was stored in a freezer and, immediately before the HS-AFM experiment, thawed and dissolved in a 10 mM $MgCl_2$ solution at a final concentration of 0.2 mg $mL^{-1}$. The mixture was sonicated for 1 min using an ultrasonic bath sonicator (AS ONE, AUC-06L) to produce small unilamellar vesicles. 2 μL of the sonicated liposome solution was deposited onto a freshly cleaved mica surface, incubated for 5 minutes, and rinsed with excess Milli-Q water (>100 μL) and 20 μL of AFM-B buffer. 1 μL of the DNA loading reaction was deposited onto the lipid membrane imaging stage and incubated for 5 min. HS-AFM imaging was performed in AFM-B buffer containing 5 mM of the indicated adenosine derivatives, after rinsing with the same buffer.

**HS-AFM data analysis**

All HS-AFM images and movies are presented after being processed with a Gaussian-kernel filter (standard deviation ~1 nm) and color scaling adjustments. The individual HS-AFM datasets consist of multiple frames; however, the offset in the Z direction varies for each frame due to scanner drift. Conventionally, color scaling has been performed based on the maximum and minimum height values.

However, this method results in an unstable color scale across frames due to the influence of noise and dust adsorption/desorption. To standardize the color scale across frames, we developed a method called PeakRef. In this approach, pixels are sorted based on their height values within each frame, and the substrate height is obtained by identifying the height value that occurs most frequently. Color scaling is then applied using the obtained substrate height, ensuring consistent color scaling across all frames. In Figs. 2A,D, 4E, and 6A, a frame averaging method with a triangular window over the previous and subsequent frames was also applied to reduce noise and enhance the static structures.

To calculate the elbow angle of Smc5/6, three control points were manually placed along the ridgeline of the molecular arms (fig. S4), and the angle between the two straight lines (vectors of **a** and **b**) connecting these points was calculated using a trigonometric equation expressed by

$$\theta = \cos^{-1}\left(\frac{\mathbf{a}\cdot\mathbf{b}}{\|\mathbf{a}\|\|\mathbf{b}\|}\right). \quad (1)$$

To evaluate the curvature distribution of DNA (fig. S8B,C), the curvature angle was calculated at each point of the DNA. To determine the pseudo-curvature angle of a smooth line, the point of interest on the curve was locally approximated as two straight lines within the range of ±30 nm, a value slightly shorter than the persistence length of DNA (~50 nm), and the angle between these straight lines was calculated.

To quantify the compactness of DNA (Fig. 6B,C), we calculated the radius of gyration ($R_{\mathrm{G}}$), which is expressed as

$$R_{\mathrm{G}} = \sqrt{\frac{1}{N}\sum_{i=1}^{N}\left(\mathbf{r}_i - \mathbf{r}_{\mathrm{CM}}\right)^2}, \quad (2)$$

where $\mathbf{r}_{\mathrm{CM}}$ represents the coordination vectors of the center of mass, and $N$ denotes the total number of points. Initially, several tens of control points were manually placed along the DNA contour. These points were then interpolated using a spline method to increase the total number of points to 800. Given that the total length of the DNA was ~644 nm, this corresponds to a model of beads strung together at intervals of ~0.8 nm. For the calculation of $R_G$, we used data acquired within the first ten frames to minimize the effect of cantilever scanning.

**Prediction of the ATP-dependent conformational change of Smc5/6**

We performed a molecular simulation using a variant of the freely-jointed chain (FJC) model (*43*), with data assimilation from our HS-AFM results (Fig. 2) and the published cryo-EM structures of Smc5/6, to predict the molecular trajectory of ATP-dependent conformational changes of the Smc5/6 hexamer (fig. S3, movie S1). The simulation and atomic model visualization were performed with lab-built software utilizing the OpenGL library (Khronos Group, https://www.opengl.org/sdk/).

First, a coarse-grained (CG) model was constructed based on the all-atom structural model of the apo state Smc5/6 hexamer (I-form) derived from the cryo-EM structure (PDB-7QCD) (*25*). In this model, CG-particles, denoted as 'p', were connected by chains of rigid segments (fig. S3). Here, $p_n$ represents the *n*-th CG-particle, corresponding to the Cα of amino acids identified by their residue sequence numbers, as listed in table S1. The 10 CG particles were placed at the following amino-acid residues: E187 (p1) and T200 (p2) on the Smc5 ATPase head, D739 (p3) on the Smc5 arm, Q647 (p4) and N618 (p5) on the Smc5 head, N503 (p6) on the Smc6 hinge, V761 (p7) and I805 (p8) on the Smc6 arm, and T1035 (p9) and S229 (p10) on the Smc6 ATPase. These 10 CG-particles were connected by 9 segments with fixed distances, which were maintained according to the initial structure from PDB-7QCD to avoid changes during the simulation. The 9 segments were further classified into 6 parts, denoted as 'S' (table S2): S1 (p1-p2-p3), S2 (p3-p4), S3 (p4-p5-p6), S4 (p6-p7), S5 (p7-p8) and S6 (p8-p9-p10). The angles between CG-particles within S1, S3, and S6 were fixed at their initial values, while the angles between the parts were allowed to vary arbitrarily.

This CG framework model was converted into the O-form by pulling p3 and p7 in opposite directions. During this process, a harmonic oscillator model-type force, which mimics ATP-dependent head engagement, was applied between p1 and p10 to maintain p1–p10 distance at 1.2 nm. Finally, an all-atom model of the O-form was generated by projecting the all-atom model of the I-form onto the coordination of the CG model of the O-form for each part.

Pseudo-AFM images were generated with lab-built software implementing a tip-convolution algorithm inspired by that used in the SPM simulator (*59*, *60*). An ideal symmetrical hemisphere with a tip radius of 5 nm, terminating in a cone-shaped tip with a half-angle of 10 degrees, was assumed. The tip was scanned by tracing the topmost surface of the molecular structure, and the tip trajectory was reconstructed into a pseudo-AFM image. Molecules were treated as rigid, and their stiffness, fluctuations, and scanning feedback errors were not considered.

**Figures S**

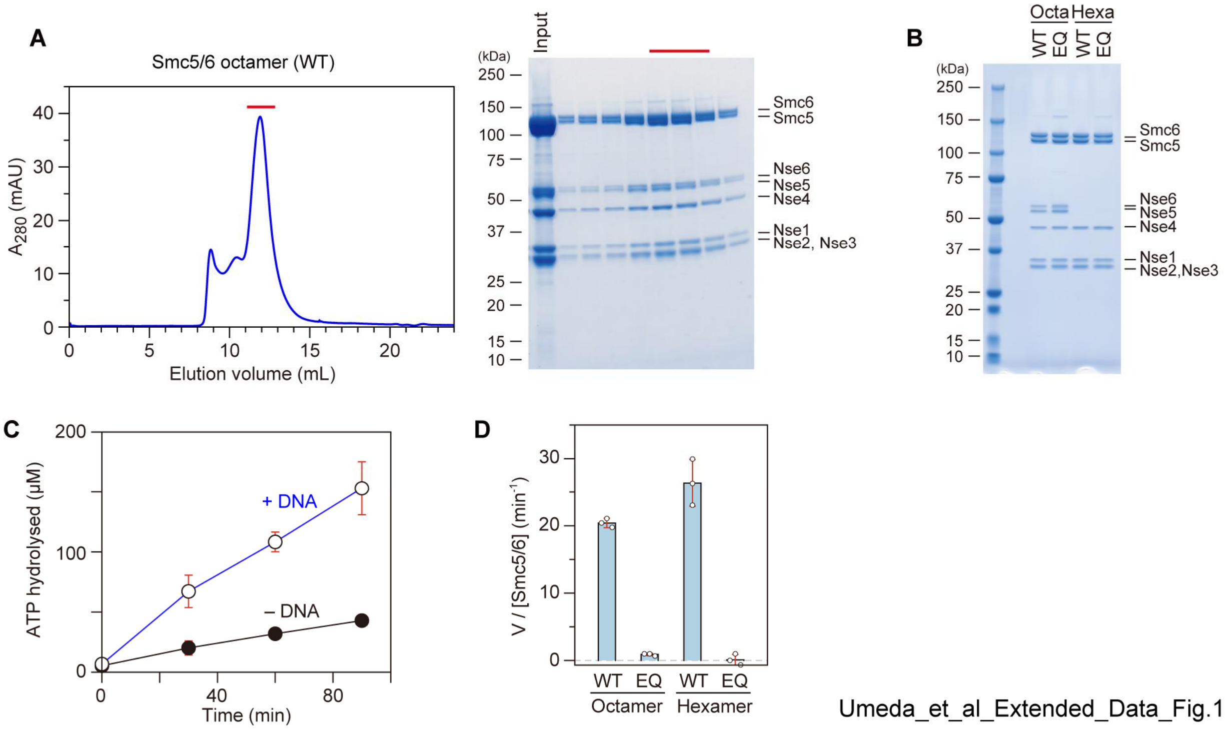


**Figure S1. Purification of the budding yeast Smc5/6 complex.**

(**A**) Size-exclusion chromatogram profile of the Smc5/6 octamer (left). The fractions were analyzed by SDS-PAGE followed by Coomassie brilliant blue (CBB) staining (right). The peak fractions, indicated by red lines, were collected and used for HS-AFM and biochemical analyses.

(**B**) Purified wild-type and EQ mutant Smc5/6 hexamers and octamers were analyzed by SDS-PAGE followed by CBB staining.

(**C**) Time-course of ATP hydrolysis by the Smc5/6 octamer with or without DNA.

(**D**) ATPase activities of the indicated Smc5/6 complexes in the presence of DNA, which were measured from the concentration of phosphate products at the 30-minute time point following the initiation of the reaction. In panels c and d, data are presented as means ± s.d. from three independent experiments.

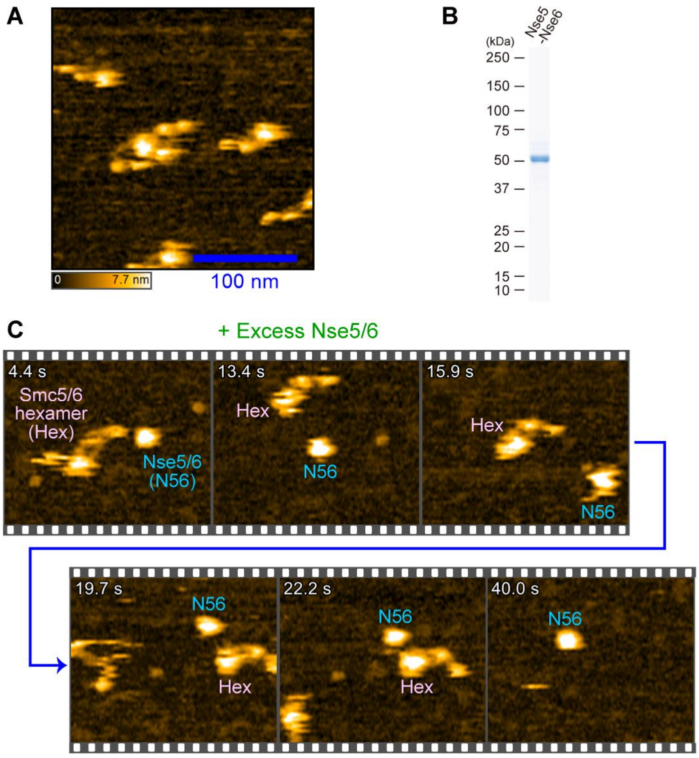


Umeda_et_al_Extended_Data_Fig.2

**Figure S2. HS-AFM observation of the Smc5/6 octamer using a mica imaging stage.**

(**A**) HS-AFM observation of the Smc5/6 octamer in a buffer containing 300 mM KOAc. The octamer consists of the Smc5/6 hexamer and the DNA-loading accessory factor, the Nse5–Nse6 heterodimer (Nse5/6). Nse5/6 were co-purified with Smc5/6 hexamer during gel filtration chromatography (fig. S1A), indicating a stable octameric complex in solution. However, after deposition on mica, we predominantly observed ‘SMC’-shaped molecules corresponding to the Smc5/6 hexamer state (Fig. 1A). Scan area, 250 × 250 $nm^2$ at 200 × 100 $pixels^2$. Imaging rate: 1.3 s per frame.

(**B**) Purification of the Nse5/6 heterodimer. Purified Nse5/6 was analyzed by SDS-PAGE followed by CBB staining.

(**C**) Nse5/6 appeared to dissociate from the Smc5/6 core complex when placed on mica. We observed the Smc5/6 octamer after the addition of excess amounts of the purified Nse5/6. Adsorption of Nse5/6 on mica was seen when 10-fold excess amounts of Nse5/6 compared to Smc5/6 octamer was applied. This suggests that Nse5/6 was not stably adsorbed on mica and was prone to detachment. Under these conditions, we occasionally observed Smc5/6 molecules corresponding to the hexamer state (Hex) and Nse5/6 (N56) in the same viewing window, but they never physically interacted with each other. These results strongly suggest that when the Smc5/6 octamer was placed on mica, the physical interaction between the Smc5/6 hexamer and Nse5/6 became unstable. Consequently, the Nse5/6 dissociated from the Smc5/6 hexamer and detached from mica, preventing visualization of the Smc5/6 octamer under the current experimental conditions. Due to this limitation, this study focused on the hexamer for the visualization of the Smc5/6 complex.

Scan area, 200 × 120 $nm^2$ at 200 × 60 $pixels^2$, cropped to 150 × 120 $nm^2$. Imaging rate: 0.63 s per frame.

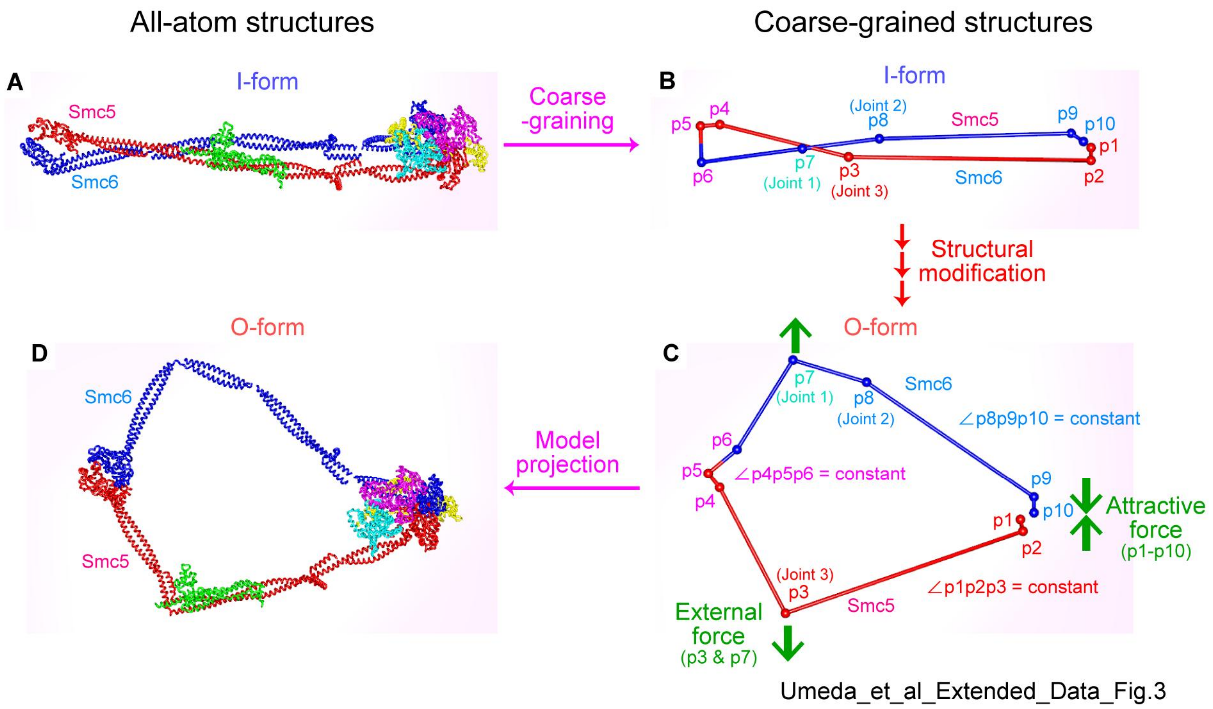


**Figure S3. Atomic model construction of the ATP-bound Smc5/6 hexamer.**

(**A**) The atomic model of apo-state Smc5/6 complex from cryo-EM structure (PDB-7QCD).

(**B**) A coarse-grained (CG) framework generated from (A). As described in Materials and Methods, we placed 10 CG-particles (denoted as p) at the following amino-acid residues, which were connected by 9 rigid chains: E187 (p1) and T200 (p2) on the Smc5 ATPase head, D739 (p3) on the Smc5 arm, Q647 (p4) and N618 (p5) on the Smc5 head, N503 (p6) on the Smc6 hinge, V761 (p7) and I805 (p8) on the Smc6 arm, and T1035 (p9) and S229 (p10) on the Smc6 ATPase (see also table S1). The ATPase head regions were defined as E187 (p1) – T200 (p2) for Smc5 and S229 (p10) – T1035 (p9) for Smc6, as these residues are almost linearly aligned in the atomic model of the ATP bound Smc5/6 hexamer derived from the cryo-EM structure (PDB-7TVE).

(**C**) Conversion of the I-form framework to the O-form. The I-form framework was converted to the O-form by applying virtual pulling forces at the middle of SMC arms (p3 and p7) in opposite directions. In this framework model, we fixed the angles of pairs of two chains defined as S1 (p1-p2-p3), S3 (p4-p5-p6) and S6 (p8-p9-p10) (table S2). A harmonic oscillator model-type force was also applied between p1 and p10 to recapitulate the ATP-dependent heads engagement, maintaining the p1–p10 distance at 1.2 nm.

(**D**) The predicted atomic structure of the ATP-bound Smc5/6 complex. The atomic model of the ATP-bound form was generated by projecting the apo-state atomic model onto the coordination of the coarse-grained model of the O-form for each part.

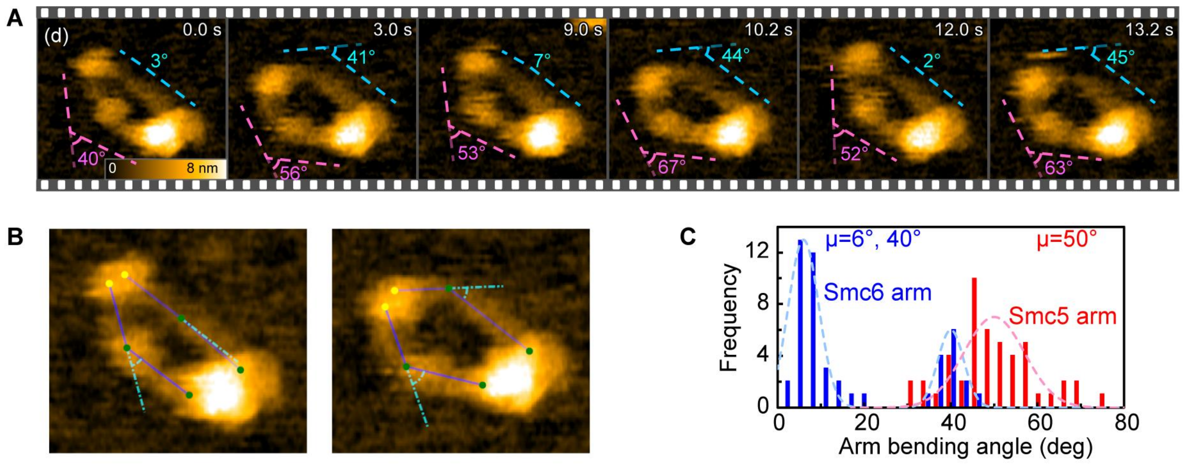


Umeda_et_al_Extended_Data_Fig.4

**Figure S4. Fluctuations of Smc5 and Smc6 arms with distinct magnitudes on mica.**

(**A**) Successive HS-AFM images of the EQ hexamer in the presence of ATP in a buffer containing 75 mM KOAc. The Smc6 arm (blue dash line) exhibited fluctuations due to repeating bending and stretching, whereas the Smc5 arm (pink dash line) remained relatively stable, maintaining a mild curvature at the Nse2 binding region. Scan area, 120 × 120 $nm^2$ at 240 × 120 $pixels^2$, cropped to 64 × 54 $nm^2$. Imaging rate: 0.6 s per frame.

(**B**) Measurement of SMC arm bending angles. For each SMC arm, ridgelines consisting of two segments were placed, connecting the hinge to the head. The yellow and green dots represent the control points of the hinge domain and other domains. The bending angle was then measured at the junction between the segments (see also Materials and Method).

(**C**) Distributions of Smc5 and Smc6 arm bending angles, based on a total of 48 frames from the successive HS-AFM images shown in (A).

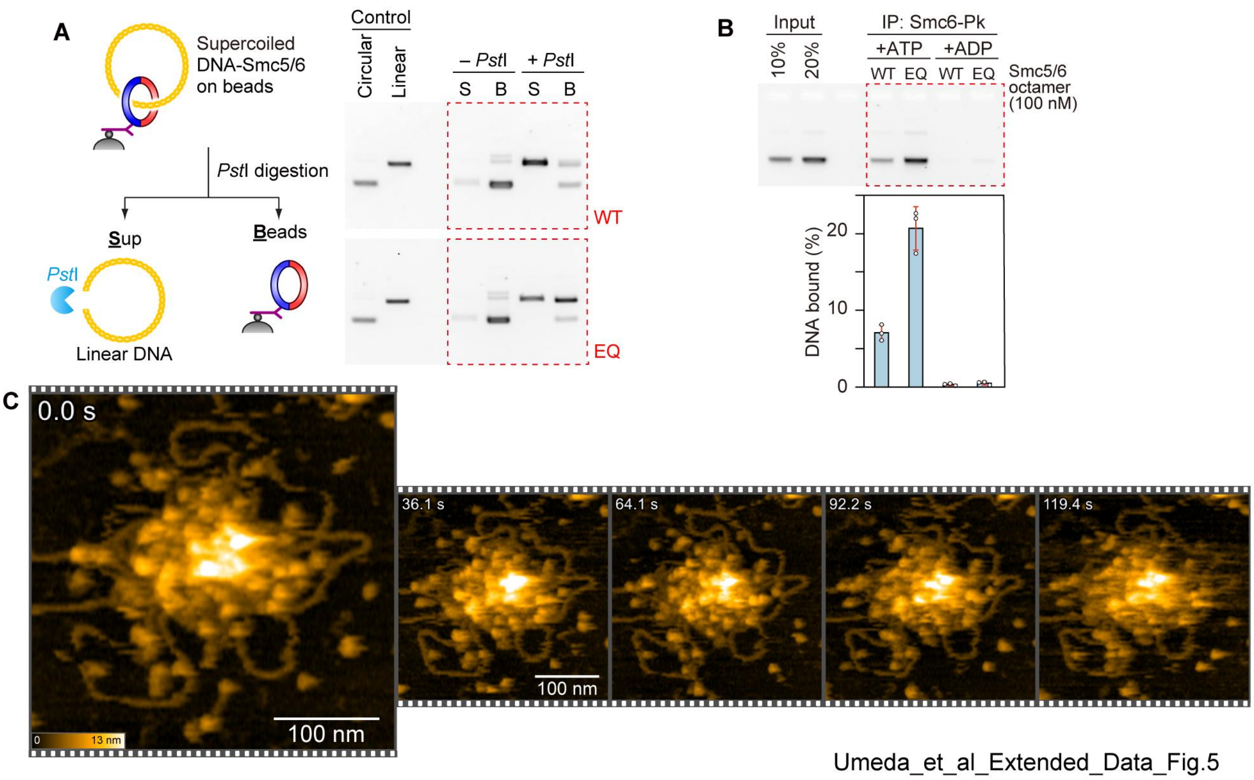


**Figure S5. Biochemical reconstitution of topological DNA entrapment by Smc5/6.**

(**A**) Schematic and gel images of the DNA-release experiments using restriction digestion to confirm the topological nature of DNA loading. Smc5/6 was initially loaded onto supercoiled plasmid DNA in the presence of ATP as Fig. 4A. After immunoprecipitation and high-salt washing to retrieve Smc5/6, the Smc5/6-bound DNA was digested with restriction enzyme *Pst*I. The cleaved DNA was released into the supernatant, while undigested DNA remained in the Smc5/6-bound bead fraction. This result indicates that linearized DNA was able to escape from Smc5/6.

(**B**) Same as Fig 4B, except that the experiment was carried out using the EQ mutant octamer complex in addition to the wild-type.

(**C**) Successive HS-AFM images of DNA-bound Smc5/6 using a mica imaging stage under a buffer with a low concentration of KOAc. Scan area, 400 × 400 $nm^2$ at 240 × 120 $pixels^2$. Imaging rate: 0.8 s per frame. Since both mica and DNA are negatively charged, DNA cannot be adsorbed on mica without modifying the surface. This can be achieved by using an imaging buffer containing potassium ions, with a salt concentration of approximately several tens of mM. Under these conditions multiple Smc5/6 molecules were found to stack on the DNA. Additionally, free Smc5/6 molecules could also be adsorbed onto the mica surface, making it difficult to distinguish DNA-bound Smc5/6 from free molecules adsorbed near the DNA. Therefore, we used the lipid membrane stage system to specifically observe DNA-bound Smc5/6 using HS-AFM.

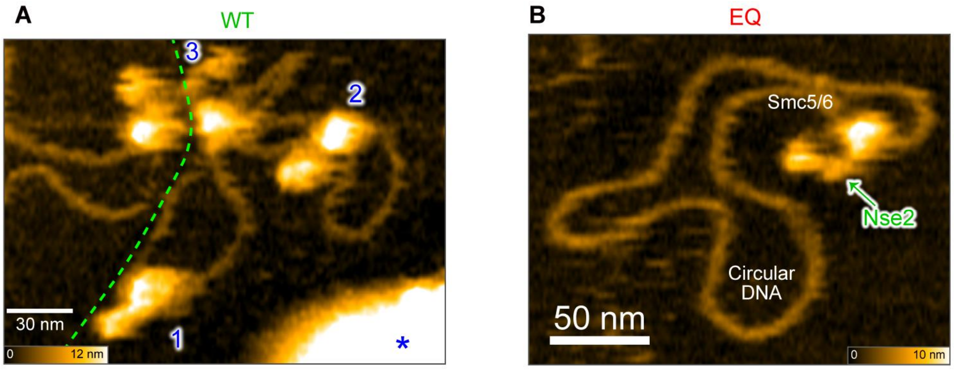


Umeda_et_al_Extended_Data_Fig.6

**Figure S6. Examples of multiple wild-type and single EQ Smc5/6 bound to DNA using a lipid membrane stage system.**

(**A**) HS-AFM images showing DNA molecules bound by multiple wild-type Smc5/6 molecules. The green dashed line indicates the boundary between the two DNA molecules, and only those loaded onto the DNA on the right are numbered. Scan area, 250 × 200 $nm^2$ at 200 × 80 $pixels^2$, cropped to 208 × 150 $nm^2$. Imaging rate: 0.7 s per frame.

(**B**) Overview scans of the DNA-bound EQ mutant Smc5/6 molecule (movie S4B). Scan area, 300 × 240 $nm^2$ at 240 × 96 $pixels^2$, cropped to 200 × 156 $nm^2$. Imaging rate, 1 s per frame.

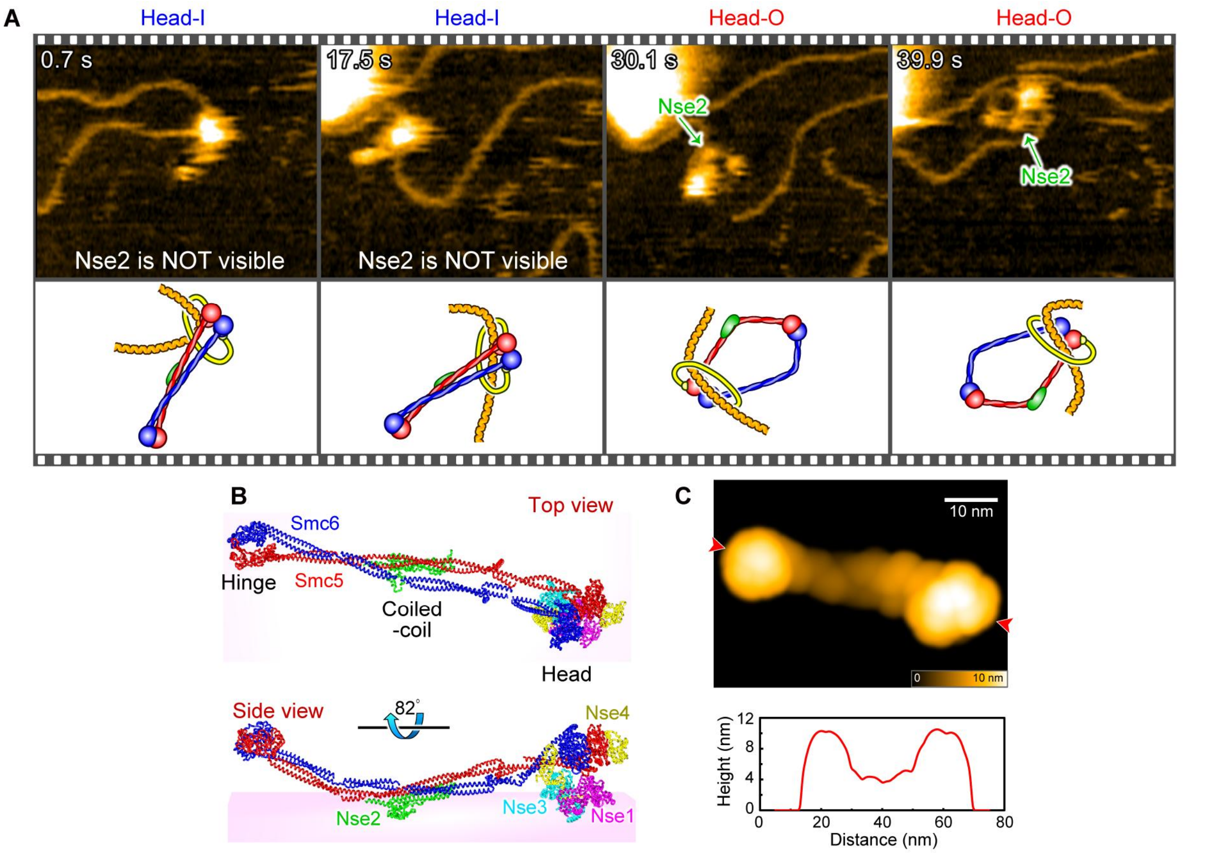


Umeda_et_al_Extended_Data_Fig.7

**Figure S7. DNA-bound Head-I Smc5/6 was adsorbed on the lipid membrane stage in an orientation distinct from that on mica.**

(**A**) HS-AFM images showing the transition of DNA-bound Smc5/6 from the I-form to the O-form. In the first two frames, DNA-bound Smc5/6 exhibited the I-form structure; however, unlike the results obtained on mica, Nse2 was not detected on the Smc5 arm. In contrast, the Nse2 signal appeared when the I-form was converted to the O-form in the 30.1 s frame.

(**B**) Atomic model of the budding yeast Smc5/6 hexamer in the apo state derived from cryo-EM structure (EMD-13895, PDB-7QCD), adsorbed onto the surface in the orientation opposite to that shown in Fig. 2B, with Nse2 facing down.

(**C**) A pseudo-AFM image of the Smc5/6 hexamer from panel **b**. In contrast to Fig 2B, Nse2 was not visible in the middle of the SMC arm region. The corresponding line profile of the molecular height across the two red arrowheads is presented. These results suggest that the DNA-bound I-form of Smc5/6 was adsorbed on the lipid membrane surface with Nse2 facing downwards, in contrast to its orientation on mica.

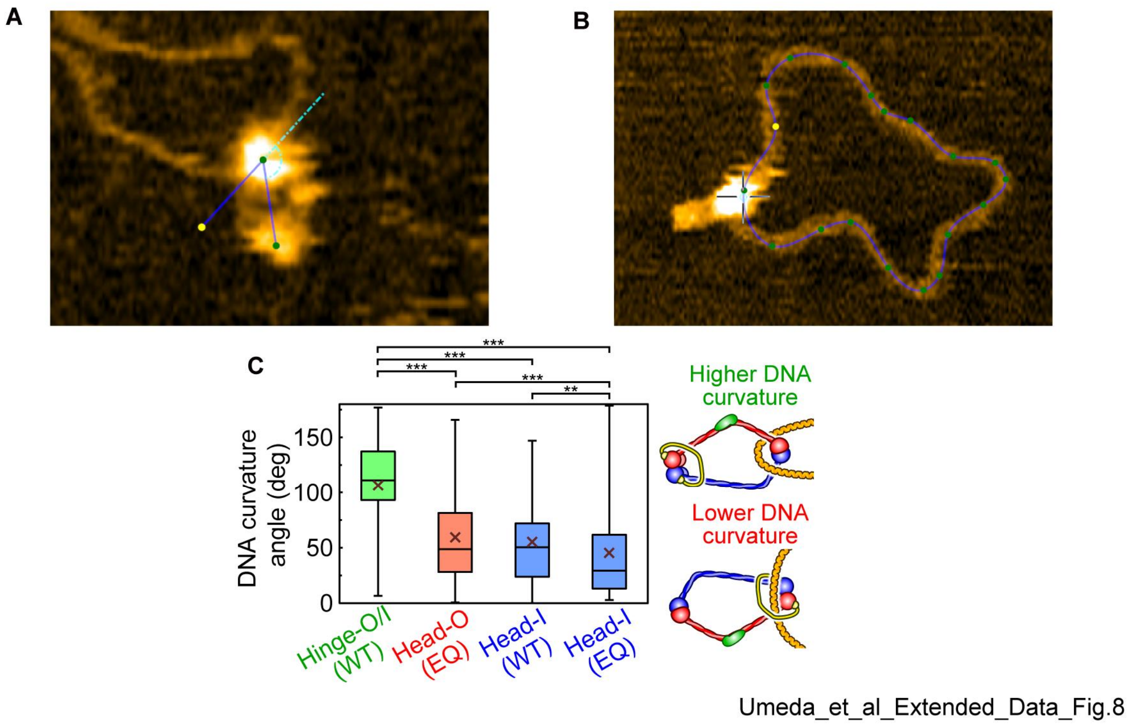


**Figure S8. Smc5/6 binding at the hinge or head exhibits distinct molecular behaviors on DNA.**

(**A**) Measurements of Smc5/6 binding angles relative to DNA (see Materials and Methods).

(**B**) Measurements of DNA curvature angles at Smc5/6 binding sites (see Materials and Methods). The cross-hair indicates the anchor point corresponding to the molecular binding site.

(**C**) Quantification of DNA curvature angles, based on a total of 165, 236, 147, and 143 frames for WT Hinge-O/I, WT Head-O, WT Head-I, and EQ Head-I, respectively ($n$ = 3 molecules per condition) from successive HS-AFM images. In the box-and-whisker plot, crosses indicate mean values. Statistical differences were assessed using the nonparametric Brunner-Munzel test (***: $p < 0.001$).

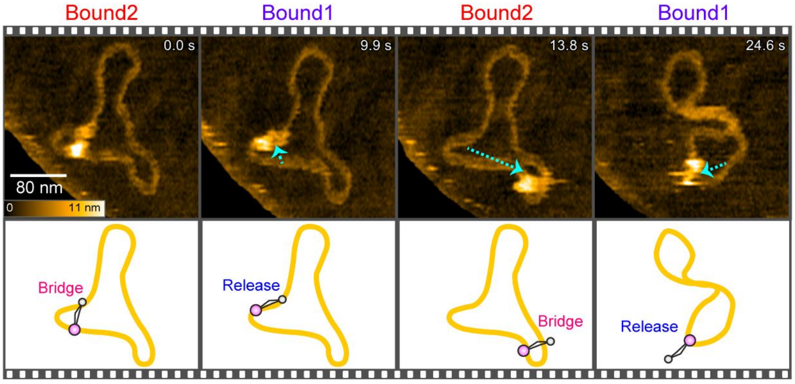


Umeda_et_al_Extended_Data_Fig.9

**Figure S9. Transient DNA bridging by Smc5/6 in the "Bound 2 (bridge)" state.**

Successive HS-AFM images of Smc5/6 bound DNA in the "Bound 2" state. Smc5/6 repeatedly bridged two DNA segments and released one of them during HS-AFM observation. Based on these results, we concluded that the DNA bridging observed in the "Bound 2" state was transient. Scan area, 350 × 280 $nm^2$ at 200 × 80 $pixels^2$, cropped to 280 × 250 $nm^2$. Imaging rate: 0.98 s per frame.

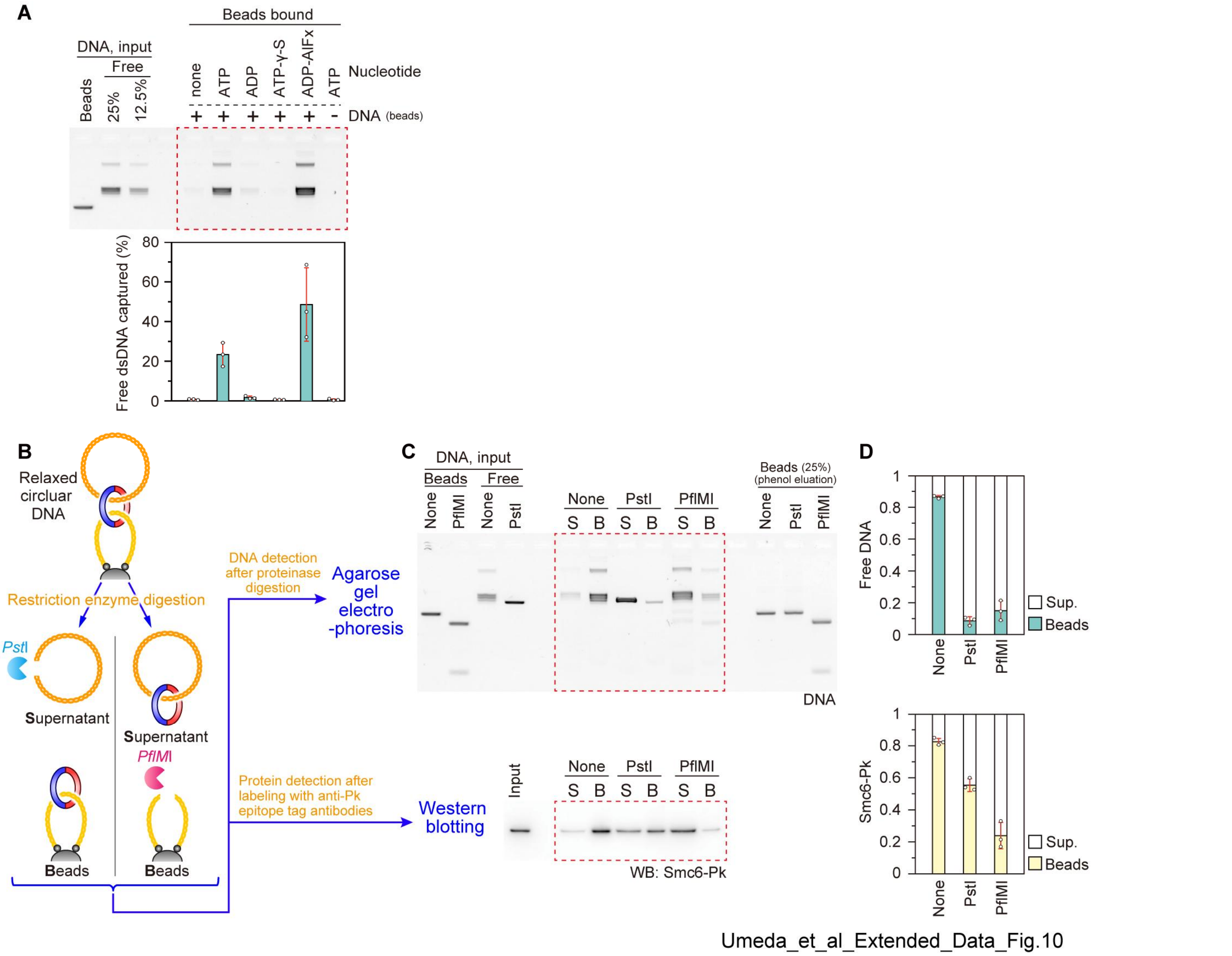


**Figure S10| Smc5/6 mediates DNA–DNA tethering through topological DNA entrapment.**

(**A**) Representative agarose gel image of the DNA–DNA tethering assay in the absence or presence of the indicated nucleotide derivatives. The graph shows the quantification of captured free DNA (obtained from three independent experiments; mean ± s.d.).

(**B**) Schematic of DNA-release experiments mediated by restriction digestion. After retrieving DNA–DNA tethering products as depicted in Fig. 6D, captured free DNA and DNA immobilized on the magnetic beads was digested by *Pst*I and *Pfl*MI restriction enzymes, respectively. The captured free DNA is expected to be released into the supernatant if DNA–DNA tethering requires topological entrapment. Likewise, digestion of bead-bound DNA would also release Smc5/6 into the supernatant. Afterward, electrophoresis and western blotting were performed.

(**C**) Representative agarose gel (top) showing free DNA tethered by Smc5/6 to immobilized DNA, and western blot (bottom) detecting DNA-bound Smc5/6, described in panel **b**.

(**D**) Quantification of captured free DNA (top) and Smc5/6 bound to DNA beads (bottom) ($n$ = 3 independent experiments; mean ± s.d.). The majority of captured free DNA was released from immobilized DNA beads by digestion of free or bead-bound DNA. In addition, a significantly larger amount of Smc5/6 was released into the supernatant upon linearization of bead-bound DNA. These results strongly suggest that Smc5/6 mediates DNA–DNA tethering.

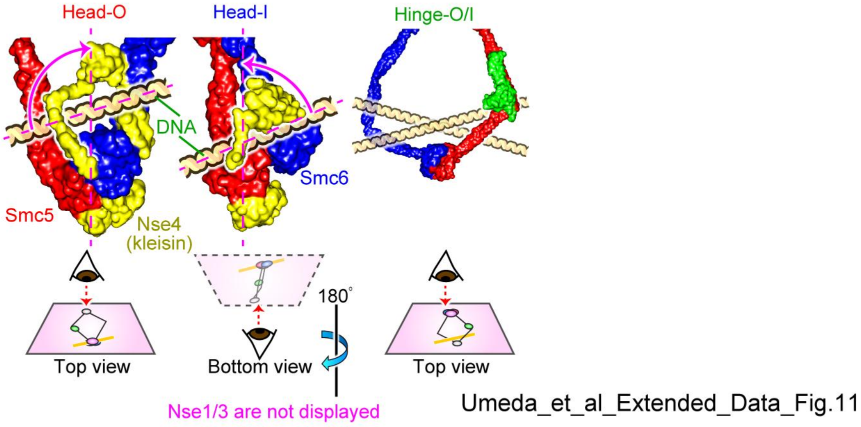


**Figure S11. Predicted molecular models of Smc5/6 that topologically entrap DNA.**

Structural models of the DNA entrapment state of Smc5/6. Left: Head-O, middle: Head-I, right: Hinge-O/I. In the Head-O and Head-I models, DNA strands were positioned within the Nse4 kleisin compartment at the tilt angles observed in our HS-AFM experiments (Fig. 5F), and then fitted into the tilted gap between the kleisin and head domains. Motion of DNA within the Nse4 kleisin appears to be constrained due to the narrower space in the Head-I state, compared to the Hinge-O/I state.

**Supplementary Table**

Table S1 Assignment of atomic numbers to beads for FJC model construction.

| CG-Particle Name | Residue Sequence Number | Atom Serial Number in PDB-7QCD | Subdomain Name | Function Name |
|---|---|---|---|---|
| p1 | E187 | 1202 | Smc5 | ABC Domain |
| p2 | T200 | 1309 | Smc5 | Head |
| p3 | D739 | 5649 | Smc5 | Elbow |
| p4 | Q647 | 4883 | Smc5 | Hinge |
| p5 | N618 | 4647 | Smc5 | Hinge |
| p6 | N503 | 11495 | Smc6 | Hinge |
| p7 | V761 | 13610 | Smc6 | Elbow |
| p8 | I805 | 13951 | Smc6 | Elbow |
| p9 | T1035 | 15728 | Smc6 | Head |
| p10 | S229 | 9453 | Smc6 | ABC Domain |

Table S2 Assignment of residue numbers to parts for FJC model construction.

| Part Name | CG-Particle Range | Residue Sequence Range | Subdomain Name |
|---|---|---|---|
| S1 | p1-p2-p3 | 1–369 | Smc5 |
| | | 738–1068 | Smc5 |
| | | 1–267 | Nse2 |
| | | 1–336 | Nse1 |
| | | 1–303 | Nse3 |
| | | 125–402 | Nse4 |
| S2 | p3-p4 | 370–450 | Smc5 |
| | | 648–737 | Smc5 |
| S3 | p4-p5-p6 | 451–647 | Smc5 |
| | | 503–692 | Smc6 |
| S4 | p6-p7 | 436–502 | Smc6 |
| | | 693–759 | Smc6 |
| S5 | p7-p8 | 371–435 | Smc6 |
| | | 760–813 | Smc6 |
| S6 | p8-p9-p10 | 1–370 | Smc6 |
| | | 814–1104 | Smc6 |
| | | 1–124 | Nse4 |

**Movie S legends**

**Movie S1.** Simulated molecular transition of the Smc5/6 hexamer from the I-form to O-form via an external force, related to Fig. 2E.

**Movie S2.** (**A-C**), HS-AFM videos of the wild-type Smc5/6 hexamer in the absence of ATP (A), in the presence of ATP (B), and of the EQ mutant in the presence of ATP (D), observed on the mica imaging stage (related to Fig. 3A).

(**A**): Scan area, 100 × 100 $nm^2$ at 160 × 80 $pixels^2$, cropped to 90 × 86 $nm^2$.

(**B**): Scan area, 150 × 150 $nm^2$ at 160 × 80 $pixels^2$, cropped to 110 × 105 $nm^2$.

(**C**): Scan area, 100 × 100 $nm^2$ at 160 × 80 $pixels^2$, cropped to 95 × 90 $nm^2$.

Imaging rate: 0.5 s per frame for (A) and (C), 0.7 s per frame for (B).

**Movie S3.** HS-AFM videos of the wild-type Smc5/6 hexamer showing the collapsed conformation upon Nse2 dissociation, observed on the mica imaging stage, related to Fig. 3D.

Scan area, 100 × 100 $nm^2$ at 200 × 100 $pixels^2$, cropped to 81 × 70 $nm^2$.

Imaging rate, 0.3 s per frame.

**Movie S4|** (**A**,**B**) Overview HS-AFM videos of DNA-bound wild-type (A) and EQ mutant (B) Smc5/6, observed on the lipid membrane imaging stage (related to Fig. 4E and fig. S6B).

(**A**): Scan area, 350 × 280 $nm^2$ at 200 × 80 $pixels^2$, cropped to 298 × 238 $nm^2$.

(**B**): Scan area, 300 × 240 $nm^2$ at 240 × 96 $pixels^2$, cropped to 255 × 204 $nm^2$.

Imaging rate for both a and b: 1 s per frame.

**Movie S5.** HS-AFM video of two EQ mutant Smc5/6 complexes bound to a single DNA molecule, observed on the lipid membrane imaging stage (related to Fig. 4F).

Scan area, 220 × 176 $nm^2$ at 200 × 80 $pixels^2$, cropped to 187 × 134 $nm^2$.

Imaging rate: 0.6 s per frame.

**Movie S6.** (**A-C**), HS-AFM videos of DNA-bound wild-type Smc5/6 in the Hinge-O/I state (A), in the Head-I state (B), and of DNA-bound EQ mutant Smc5/6 in the Head-O state (C), observed on the lipid membrane imaging stage (related to Fig. 5B–D).

(**A**): Scan area, 150 × 150 $nm^2$ at 160 × 80 $pixels^2$, cropped to 150 × 110 $nm^2$.

(**B**): Scan area, 150 × 120 $nm^2$ at 160 × 64 $pixels^2$.

(**C**): Scan area, 220 × 176 $nm^2$ at 200 × 80 $pixels^2$, cropped to 172 × 125 $nm^2$.

Imaging rates: 0.37 s per frame (A), 0.34 s per frame (B), 0.6 s per frame (C).
